# Gcn5l1 governs heart-to-liver crosstalk in cardiometabolic heart failure by modulating cholesterol metabolism

**DOI:** 10.1101/2025.02.05.636634

**Authors:** Paramesha Bugga, Bellina A.S. Mushala, Michael W. Stoner, Janet R. Manning, Nisha Bhattarai, Maryam Sharifi-Sanjani, Manling Zhang, Amber Vandevender, Raja G.R. Mooli, Sadeesh K. Ramakrishnan, John C. Sembrat, Brett A. Kaufman, Sruti S. Shiva, Cassandra L. Happe, Steven J. Mullet, Stacy L. Gelhaus, Michael J. Jurczak, Iain Scott

## Abstract

Cardiometabolic diseases, such as obesity and hypertension, negatively impact several organs simultaneously [1]. These adverse outcomes may result from a single stress that disrupts similar metabolic processes in different tissues, or from specific signals that communicate damage from one organ to another. Recent systems biology studies in cardiometabolic heart failure with preserved ejection fraction (HFpEF) have identified unidirectional liver-to-heart signaling pathways that modulate cardiac inflammation and fibrosis [2, 3]. These pathways are controlled by liver-secreted factors that govern cardiac disease progression, supporting a growing paradigm whereby liver dysfunction drives disease in the heart [4]. Despite increased recognition of the heart as a secretory organ [5], its capacity to regulate metabolic processes in other tissues remains uncertain. Here, we identify Gcn5l1 as a heart-specific factor required to protect from systemic metabolic dysfunction in cardiometabolic HFpEF. Along with exacerbated cardiac dysfunction, cardiomyocyte-restricted Gcn5l1 deletion promoted obesity, systemic insulin resistance, and hepatic lipid accumulation in a mouse model of HFpEF. Integrated multiomic analyses showed that deletion of Gcn5l1 in cardiomyocytes disrupted Angptl3-regulated lipase activity across heart and liver, resulting in cholesterol-mediated liver steatosis. These changes were blocked by inhibiting Angptl3 using a clinically-approved monoclonal antibody. Our results demonstrate that heart-to-liver crosstalk pathways exist in HFpEF, and identify the heart as a source of systemic pathological signaling in cardiometabolic disease.

**Graphical abstract:** 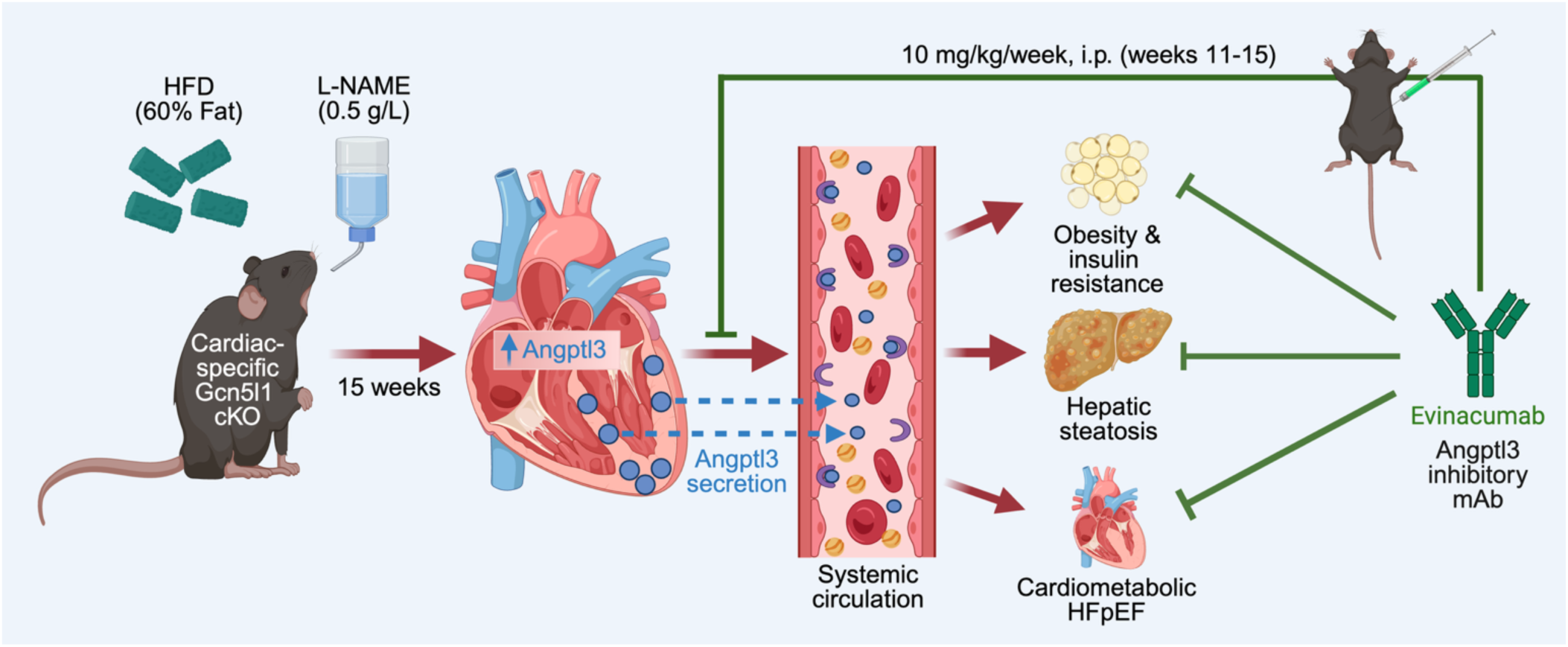

## Introduction

The prevalence of cardiometabolic heart failure with preserved ejection fraction (HFpEF) continues to grow, representing over half of the heart failure cases in the United States [6]. No specific medication for HFpEF currently exists, therefore treatment guidelines focus on the management of comorbidities related to metabolic syndrome (*e.g.* obesity, type-2 diabetes, hypertension) that promote HFpEF progression [7]. These same comorbidities also drive dysfunction in several non-cardiac tissues, and the links between cardiometabolic disease presentations in different organs are increasingly being recognized [1]. Recent systems biology studies on the potential crosstalk between HFpEF and metabolic disease in the liver (*e.g.* metabolic dysfunction-associated liver disease; MASLD) have focused on how liver dysfunction can subsequently promote disease in the heart, through the release of secreted liver proteins that control cardiac inflammation and fibrosis [2, 3]. This focus on liver-to-heart signaling may reflect the situation in the clinic, where current MASLD is a known risk factor for future HFpEF development [8]. Here, in direct contrast to this developing paradigm of liver-initiated metabolic heart disease, we report for the first time a genetic defect in the heart which drives systemic metabolic dysfunction and MASLD in the context of HFpEF.

## Results

### Cardiac Gcn5l1 governs systemic metabolism

General control of amino acid synthesis 5-like 1 (Gcn5l1; also known as Bloc1s1 and Blos1) was previously shown to regulate fuel metabolism in a number of high energy demand tissues, including the heart, liver, and kidney [9, 10, 11]. To determine its potential role in cardiometabolic disease, we placed tamoxifen-inducible, cardiomyocyte-specific Gcn5l1 knockout mice (Gcn5l1 cKO; **Supplementary Data Fig. 1a**) and wildtype controls (WT) on either a chow diet or a preclinical HFpEF diet (60% high fat diet + 0.5 g/L L-NAME in drinking water [12]) for 15 weeks (**Fig. 1a**). Unexpectedly, both male and female Gcn5l1 cKO mice on the HFpEF diet developed exacerbated obesity and systemic insulin resistance compared to WT HFpEF animals (**Fig. 1b,c** and **Supplementary Data Figs. 3a,b,q**). Both sexes of Gcn5l1 cKO mice developed HFpEF-related cardiac hypertrophy, while only male mice displayed elevated lung weights consistent with heart failure-related pulmonary edema (**Fig. 1d** and **Supplementary Data Fig. 3c,e**). Similarly, male Gcn5l1 cKO mice on a HFpEF diet displayed increased cardiac interstitial fibrosis and diastolic dysfunction (as shown by increased E/e’ and E/A ratios, and prolonged IVRT) relative to WT mice under the same conditions (**Fig. 1e,f** and **Supplementary Data Fig. 2j**). These changes were specifically related to the HFpEF diet, and were not observed with high fat diet alone (**Supplementary Data Fig. 4**). Finally, Gcn5l1 cKO mice displayed dysregulated expression of multiple genes related to cardiac muscle contraction (*e.g. Atp2a2, Ryr2, Bag3*) in bulk RNA-seq from HFpEF hearts (**Fig. 1g**). Combined, these data demonstrate that cardiomyocyte-restricted loss of Gcn5l1 expression has deleterious impacts on both intrinsic cardiac function and whole-body metabolic homeostasis.

**Fig. 1:**
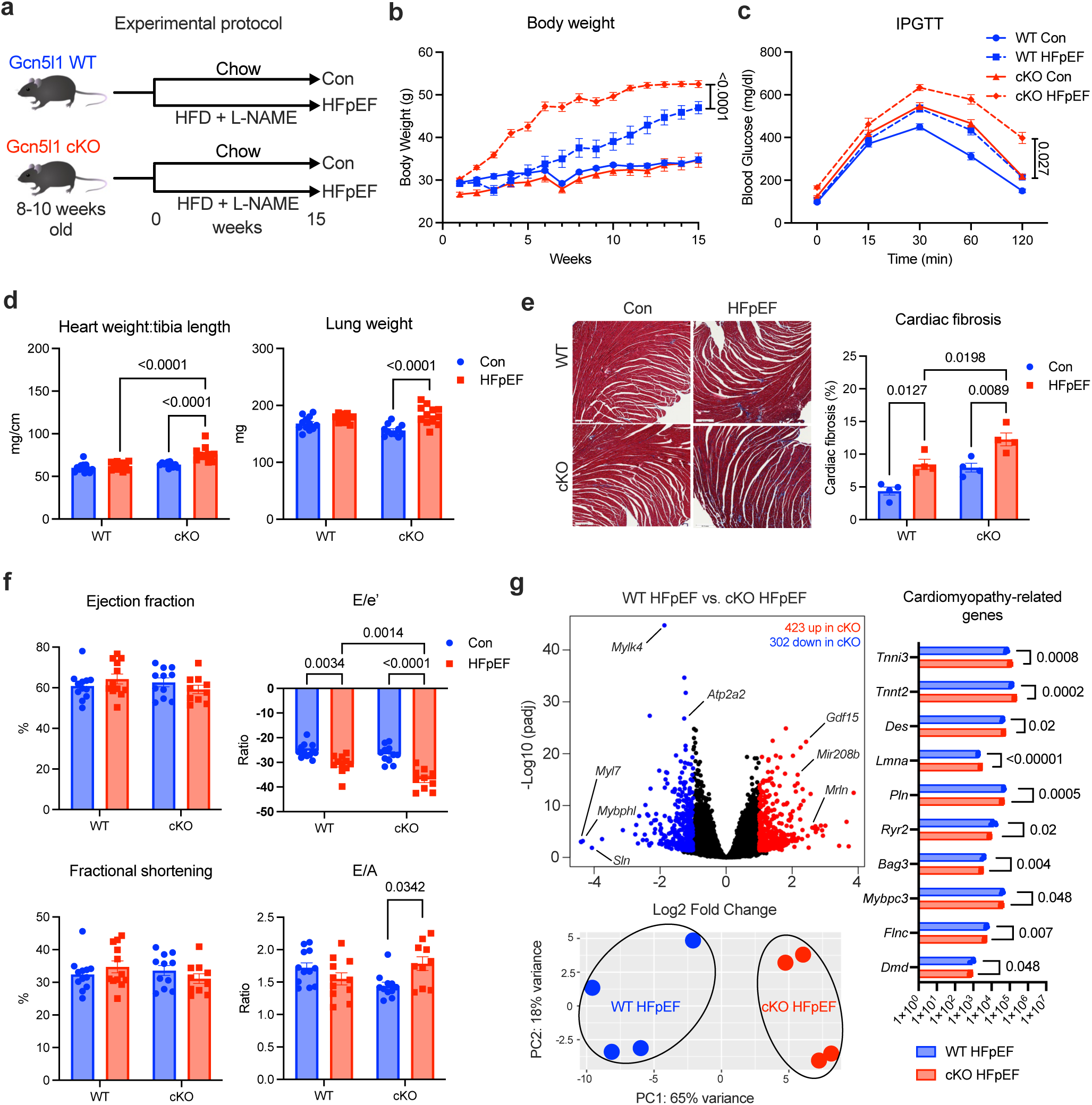
Cardiomyocyte-specific deletion of GCN5L1 under HFpEF conditions. **a,** Schematic of study protocol. **b,c,** Body weight and intraperitoneal glucose tolerance test (IPGTT) curves for WT and Gcn5l1 cKO mice on chow (Con) or 60% high fat diet + 0.5 g/L L-NAME (HFpEF) diets. Data are mean ± SEM, with two-way ANOVA at final marked time point and Tukey’s post-hoc test, N = 8. **d,** Heart and lung weights in Con and HFpEF mice. Data are two-way ANOVA and Tukey’s post-hoc test, N = 11 (WT) or 12 (cKO). **e,** Masson’s trichrome staining of cardiac fibrosis. Data are mean ± SEM, with two-way ANOVA and Tukey’s post-hoc test, N = 4. Scale bar is 200 μm. **f,** Ultrasonography analysis of cardiac systolic (ejection fraction and fractional shortening) and diastolic (E/e’ ratio and E/A ratio) function. Data are mean ± SEM, with two-way ANOVA and Tukey’s post-hoc test, N = 11 (WT) or 12 (cKO). **g,** Analysis of bulk RNA-seq from HFpEF hearts. Bar graph data are mean ± SEM, with multiple unpaired two-tailed t-tests, N = 4.

### Cardiac Gcn5l1 protects from liver steatosis

The unexpected increase in body weight and insulin resistance in Gcn5l1 cKO mice under HFpEF conditions (**Fig. 1b,c**) suggested a systemic effect of cardiomyocyte Gcn5l1 expression that extends beyond the heart. Necropsy analysis focused on the livers of Gcn5l1 cKO HFpEF mice, which displayed a ∼two-fold increase in size compared to WT controls (**Fig. 2a** and **Supplementary Data Fig. 5a,b**). Histological analysis using the NASH Activity Score (NAS) system demonstrated a significant increase in steatosis-related tissue damage in Gcn5l1 cKO HFpEF livers (**Fig. 2b,c**), while thin layer chromatography demonstrated a significant increase in liver lipids in the same organ (**Fig. 2d**). These findings suggested that a novel lipid signaling pathway, possibly originating from the heart, may be driving liver disease in Gcn5l1 cKO HFpEF mice.

**Fig. 2:**
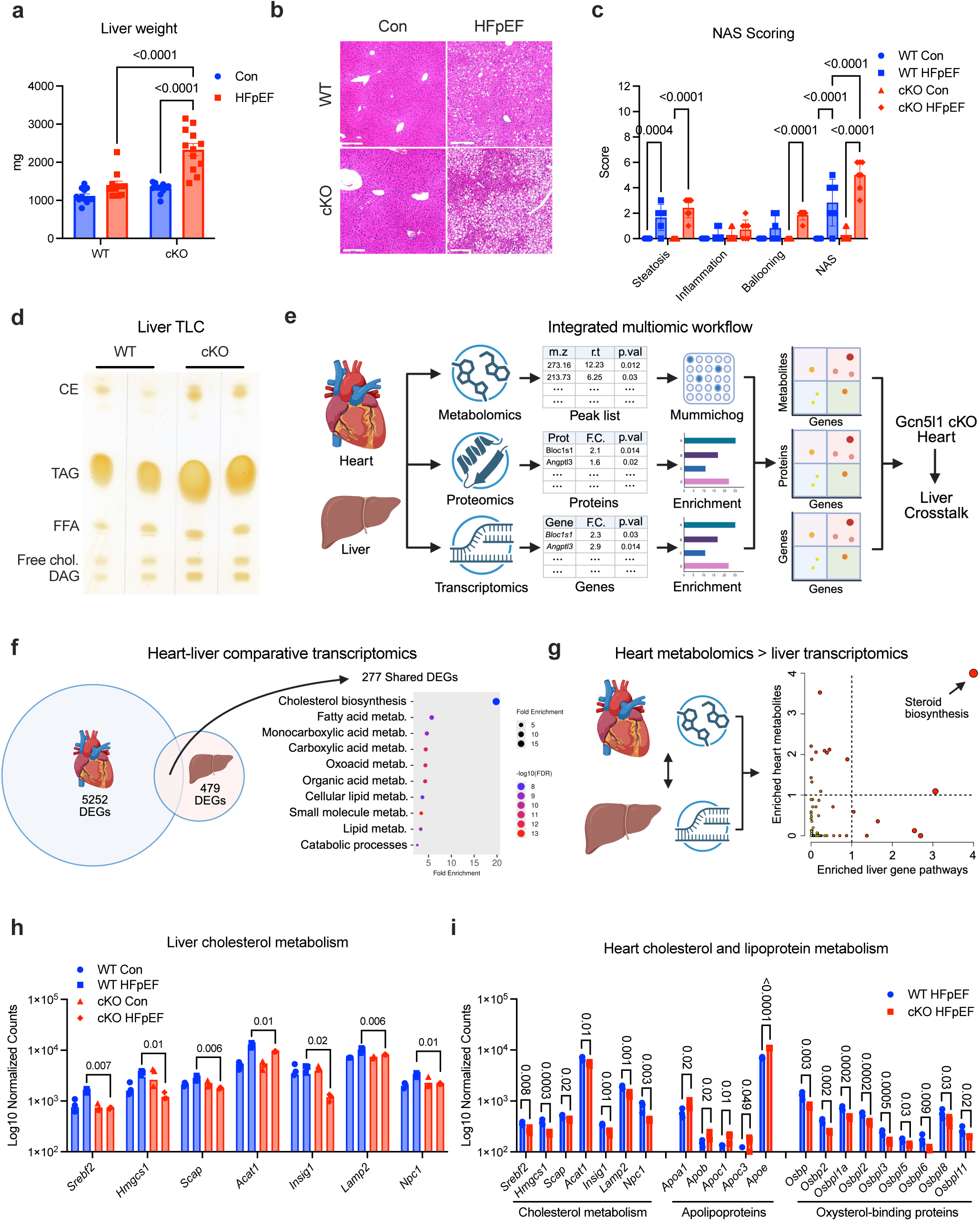
Cholesterol metabolism in Gcn5l1 cKO liver during HFpEF. **a,** Liver weights in Con and HFpEF mice. Data are mean ± SEM, with two-way ANOVA and Tukey’s post-hoc test, N = 11 (WT) or 12 (cKO). **b,c,** H&E staining and NASH Activity Score (NAS) of mouse livers. Data are mean ± SEM, with two-way ANOVA and Tukey’s post-hoc test, N = 6 (WT) or 5 (cKO). Scale bar is 200 μm. **d,** Thin layer chromatography of HFpEF liver lipid content. **e,** Diagram of integrated multiomic workflow (IMW) combining metabolomics, proteomics, and bulk RNA-seq transcriptomics to interrogate heart-to-liver crosstalk in Gcn5l1 cKO HFpEF mice. **f,** Comparative transcriptomics in heart and liver from Gcn5l1 cKO HFpEF mice vs. WT HFpEF mice. Pathway analysis performed using ShinyGo 0.85. N = 4. **g,** Joint pathway analysis of highly enriched heart metabolites and liver transcripts from Gcn5l1 cKO HFpEF mice vs. WT mice using Metaboanalyst. N = 4 (transcriptomics) or 8 (metabolomics). **h,** Liver cholesterol metabolism gene expression from bulk RNA-seq. Data are mean ± SEM, with two-way ANOVA and Tukey’s post-hoc test, N = 4. **i,** Cholesterol metabolism, apolipoprotein, and oxysterol-binding protein gene expression from bulk RNA-seq. Data are mean ± SEM, with multiple unpaired two-tailed t-tests, N = 4.

To determine the composition of this pathway, we developed an integrated multiomic workflow (IMW) that combined transcriptomic, metabolomic, and proteomic information from tissues in WT and Gcn5l1 cKO mice (**Fig. 2e**). First, we performed comparative transcriptomics from the hearts and livers of WT and Gcn5l1 cKO HFpEF animals, and found 277 differentially expressed genes (DEGs) that were shared in both organs. Pathway analysis of these DEGs showed that genes related to cholesterol biosynthesis were the most highly enriched in Gcn5l1 cKO HFpEF mice relative to WT animals under the same conditions (**Fig. 2f**). A number of these DEGs (*e.g. SCAP, NCP1, LAMP2*) were also shared in myocardial biopsies from human HFpEF patients in a recently published database (**Supplementary Data Fig. 5f** and [14]), indicating that the downregulation of cardiac cholesterol regulatory genes is a common feature in both human HFpEF and mouse models of the disease.

We next examined whether there were core metabolic features shared between the heart and liver, working under the hypothesis that cardiomyocyte-specific deletion of Gcn5l1 would drive a heart-to-liver signaling pathway. Joint pathway analysis of heart metabolites and liver transcripts from HFpEF mice demonstrated that steroid biosynthesis pathways (including cholesterol synthesis) were the most highly enriched between these tissues (**Fig. 2g**), suggesting a shared pressure on this pathway in both organs. Furthermore, we observed a significant decrease in specific metabolites from the Bloch pathway of cholesterol biosynthesis in Gcn5l1 cKO HFpEF hearts, consistent with altered cardiac cholesterol production (**Supplementary Data Fig. 5d,e**).

Targeted analysis of liver transcripts showed that the observed downregulation of cholesterol biosynthesis-related genes in Gcn5l1 cKO livers was specific to HFpEF conditions, and did not occur in chow-fed animals (**Fig. 2h**). This tallied with a lack of difference observed in gross liver morphology and histological analysis from chow-fed WT and Gcn5l1 cKO mice (**Fig. 2a-c** and **Supplementary Data Fig. 5a,b**). Biochemical analysis of livers from HFpEF mice demonstrated that Gcn5l1 cKO animals had a significant increase in cholesterol relative to WT mice (**Supplementary Data Fig. 5c**), confirming that changes to cholesterol regulatory pathways were a major component of the observed metabolic dysfunction. Combined, these data suggest that the steatosis phenotype observed in Gcn5l1 cKO HFpEF livers is due to a specific response to external HFpEF stimuli acting synergistically with Gcn5l1 depletion, as opposed to a general effect caused by Gcn5l1 deletion in the heart alone.

Finally, bulk RNA-seq analysis showed a significant decrease in multiple cholesterol metabolism genes in Gcn5l1 cKO hearts (**Fig. 2i**) that matched the pattern observed in Gcn5l1 cKO livers (**Fig. 2g**), along with pathway-wide changes in apolipoprotein and oxysterol-binding protein gene expression (**Fig. 2i**). Together, these data suggest that changes in cardiac cholesterol metabolism during HFpEF progression may stimulate metabolic disease in the liver.

### Gcn5l1 modulates a novel Angptl3-Lpl axis

We next sought to determine how changes in cholesterol metabolism may be communicated from the heart to liver in HFpEF animals. The next step in our IMW process compared heart transcriptomics and plasma proteomics, on the assumption that any heart-derived protein signaling factor would show a correlated increase in both heart transcripts and plasma protein abundance. Of the 2567 genes upregulated in Gcn5l1 cKO HFpEF hearts, there were 22 plasma proteins that showed a correlated increase in abundance (**Fig. 3a**). Of these, six proteins (Mug2, Ces3a, Ctss, Angptl3, Pigr, and Ppia) were annotated as being secreted proteins in the UniProt database, with Angptl3 being the most upregulated plasma protein between WT and Gcn5l1 cKO HFpEF mice (**Fig. 3a**). This increase was confirmed by Angptl3-specific ELISA, and was matched by a significant increase in plasma cholesterol levels (**Fig. 3b**).

**Fig. 3:**
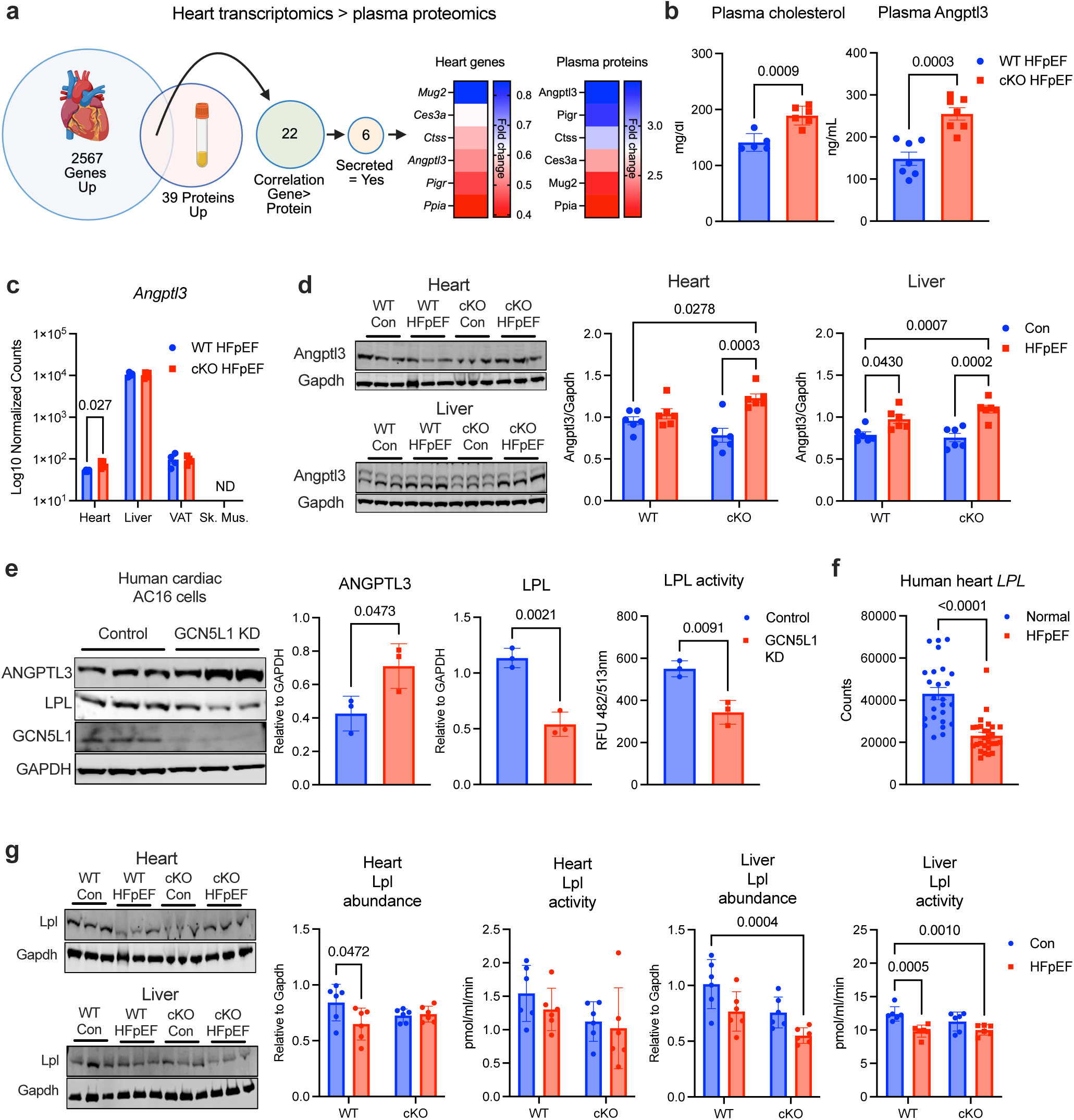
The Angptl3-Lpl axis in Gcn5l1 cKO mice during HFpEF. **a,** Identification of secreted heart signaling factors using sequential comparative analysis of heart transcriptomics and plasma proteomics from Gcn5l1 cKO HFpEF vs. WT HFpEF mice. N = 4 (transcriptomics) and 5 (proteomics; equal inputs from 5 mice pooled into a single sample for mass spectrometry analysis). **b,** Biochemical measurement of plasma cholesterol and ELISA measurement of plasma Angptl3 levels. Data are mean ± SEM, with unpaired two-tailed t-tests, N = 5 (WT cholesterol), 6 (cKO cholesterol) or 7 (WT and cKO ELISA). **c,** *Angptl3* gene expression in heart, liver, visceral adipose tissue (VAT), and skeletal muscle (Sk. Mus.) Data are mean ± SEM, with multiple unpaired two-tailed t-tests, N = 4 per tissue and genotype. **d,** Angptl3 protein abundance in heart and liver. Data are mean ± SEM, with two-way ANOVA and Tukey’s post-hoc test, N = 6. **e,** Protein abundance and Lpl activity in Control and GCN5L1 KD human cardiac AC16 cells. Data are mean ± SEM, with unpaired two-tailed t-tests, N = 3. **f,** *LPL* gene expression from a published human HFpEF data set [14]. Data are mean ± SEM, with unpaired two-tailed t-test, N = 25 (Normal) or 28 (HFpEF). **g,** Lpl protein abundance and activity in heart and liver. Data are mean ± SEM, with two-way ANOVA and Tukey’s post-hoc test, N = 6.

Angptl3 is classically viewed as a liver-secreted protein that regulates lipid abundance through the inhibition of tissue-specific lipases, such as lipoprotein lipase (Lpl) [13]. Surprisingly, analysis of several tissues demonstrated that there was a significant increase in *Angptl3* gene expression in Gcn5l1 cKO HFpEF hearts, while no increase was detected in liver, visceral adipose tissue (VAT), or skeletal muscle (Sk. Mus.; **Fig. 3c**). Western blot analysis showed higher protein abundance of Angptl3 in Gcn5l1 cKO HFpEF hearts and liver (**Fig. 3e**), which may represent either *in situ* Angptl3 production, or the binding of circulating Angptl3 to lipase targets in specific tissues. Further gene expression analyses demonstrated that there were no overall changes in cholesterol metabolism in either skeletal muscle or VAT (**Supplementary Data Fig. 6c**), suggesting that the increased Angptl3 protein abundance observed in plasma was specific to the heart-liver axis.

To resolve whether Gcn5l1 expression may regulate Angptl3 expression in the heart, we genetically depleted Gcn5l1 in human cardiac AC16 cells and rat primary neonatal cardiomyocytes. Downregulation of Gcn5l1 promoted a significant increase in ANGPTL3 protein in AC16 cells, along with a ∼two-fold increase in Angptl3 gene expression in both human and rat cardiac cell types (**Fig. 3e** and **Supplementary Data Fig. 7a,b**). Combined, these data suggest that cardiomyocytes endogenously express Angptl3 under conditions of Gcn5l1 depletion, which is upregulated further in response to cardiometabolic stress.

We then turned our attention to the *in vivo* targets of Angptl3, and examined Lpl abundance and activity in heart cells and tissues. Surprisingly, increased *ANGPTL3* gene expression and ANGPTL3 protein abundance in GCN5L1-depleted AC16 cells occurred concurrently with significant reductions in both LPL protein and activity (**Fig. 3e** and **Supplementary Data Fig. 7a**), along with decreased expression of the cholesterol metabolism proteins LDLR and NPC1 (**Supplementary Data Fig. 7c**). The observed inverse relationship between ANGPTL3 and LPL protein abundance in AC16 cells was also found in human hearts from healthy control and obese, type-2 diabetic patients (a surrogate for human HFpEF cardiac tissue; **Supplementary Data Fig. 8a**). Interrogation of a published human data set [14] demonstrated that cardiac *LPL* expression is significantly reduced in HFpEF patients (**Fig. 3f**), which was matched by a significant reduction in Lpl abundance in WT HFpEF hearts relative to chow controls (**Fig. 3g,h**).

Lipoprotein lipase is typically expressed at very low levels in adult liver; however, it can be induced under pathophysiological conditions [15]. Analysis of a published human data set [16] showed that *LPL* transcripts are significantly upregulated in patients with highly-progressed MASLD (**Supplementary Data Fig. 8b,c**), and we observed a similar increase in Gcn5l1 cKO HFpEF livers relative to WT animals (**Supplementary Data Fig. 8d**). Despite this increase in *Lpl* gene expression, we observed an inverse response at the protein and enzyme activity level in Gcn5l1 cKO HFpEF mouse livers (**Fig. 3h**). This suggests that *LPL* gene induction in MASLD patients may be a compensatory occur in response to LPL deficits at the protein level. Combined, our data suggest that loss of Gcn5l1 expression in cardiomyocytes drives atypical expression of Angptl3 in the heart in HFpEF and related cardiometabolic diseases, with subsequent remodeling of the Angptl3-Lpl axis across heart and liver.

### Angptl3 inhibition reverses dysfunction from Gcn5l1 deletion

Angptl3 is a therapeutic target in familial hypercholesterolemia, where patients with genetic deficiencies in the low-density lipoprotein (LDL) receptor pathway are resistant to cholesterol lowering strategies involving HMG-CoA-reductase inhibitors (statins) or proprotein convertase subtilisin/kexin type 9 (PCSK9) inhibitors [17]. Here, we used a recently-approved monoclonal antibody ANGPTL3 inhibitor (Evinacumab; [18]) to test whether increased abundance of Angptl3 was driving the metabolic dysfunction observed in GCN5L1 cKO HFpEF mice. WT and Gcn5l1 cKO mice were placed on a HFpEF diet for 15 weeks, with animals randomly assigned to a vehicle or Evinacumab treatment arm (10 mg/kg, once weekly via intraperitoneal injection) for the final four weeks (weeks 11 to 15; **Fig. 4a**). Evinacumab treatment successfully arrested body weight growth in Gcn5l1 cKO HFpEF mice, and significantly reduced insulin resistance in both HFpEF groups (**Fig. 4b,c**). Heart morphometry and echocardiography analyses showed that Evinacumab significantly attenuated cardiac hypertrophy and diastolic dysfunction (**Fig. 4d** and **Supplementary Data Fig. 9b**), and reduced cardiac interstitial fibrosis (**Fig. 4e**), in Gcn5l1 cKO HFpEF mice. Finally, Evinacumab treatment significantly reduced liver steatosis and organ weight in Gcn5l1 cKO HFpEF mice (**Fig. 4f**), demonstrating that the positive effects of Angptl3 inhibition in the context of HFpEF reach beyond the heart.

**Fig. 4:**
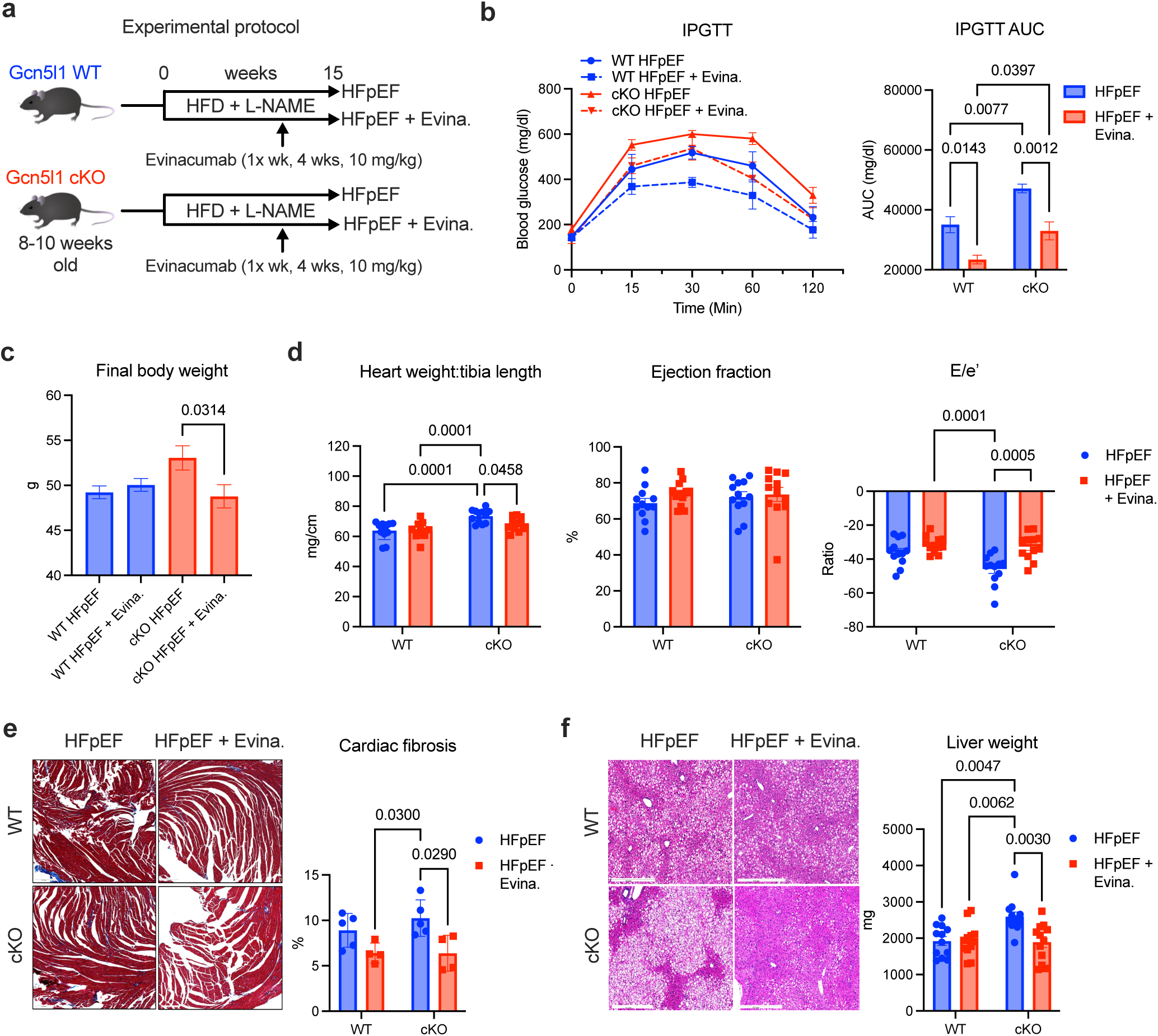
Therapeutic Angptl3 inhibition with Evinacumab during HFpEF. **a,** Schematic of study protocol. **b,** Intraperitoneal glucose tolerance test (IPGTT) and area-under-curve (AUC) analysis. Data are mean ± SEM, with two-way ANOVA and Tukey’s post-hoc test, N = 6 (WT) or 5 (cKO). **c,** Final body weight in WT and Gcn5l1 cKO mice after Evinacumab treatment. Data are mean ± SEM, with two-way ANOVA and Tukey’s post-hoc test, N = 12. **d,** Heart morphometry and ultrasonography analysis of cardiac systolic (ejection fraction) and diastolic (E/e’ ratio) function. Data are mean ± SEM, with two-way ANOVA and Tukey’s post-hoc test, N = 12. **e,** Masson’s trichrome staining of cardiac fibrosis. Data are mean ± SEM, with two-way ANOVA and Tukey’s post-hoc test, N = 5 (WT) or 4 (cKO). Scale bar is 200 μm. **f,** Liver H&E staining and organ weights. Data are mean ± SEM, with two-way ANOVA and Tukey’s post-hoc test, N = 12. Scale bar is 500 μm.

### Evinacumab rebalances the Angptl3-Lpl axis

Evinacumab treatment in humans lowers circulating LDL levels in both patients with familial hypercholesterolemia, and in healthy controls [18]. In the current study, we found that Evinacumab only reduced plasma cholesterol levels in WT HFpEF mice (**Fig. 5a**), suggesting that reduced circulating lipid levels were not the sole factor driving observed improvements in heart and liver function in Gcn5l1 cKO animals. We examined whether Evinacumab treatment may regulate cholesterol metabolism gene expression in the heart and liver, and found a widespread normalization of transcript levels in the Gcn5l1 cKO intervention group (**Fig. 5b**) compared to our initial study (**Fig. 2h,i**). We did not observe any change in cardiac cholesterol levels after Evinacumab treatment (**Fig. 5c**), but were surprised to see a significant decrease in Angptl3 transcript and protein in the hearts of Evinacumab-treated Gcn5l1 cKO HFpEF mice (**Fig. 5d** and **Supplementary Data Fig. 10a**). These changes occurred independently of any change in Lpl abundance or activity (**Fig. 5d,g**). Furthermore, Evinacumab treatment in GCN5L1-depleted AC16 cells prevented the significant increase in Angplt3 relative to control cells observed previously (**Fig. 3e** and **Supplementary Data Fig. 10c**).

**Fig. 5:**
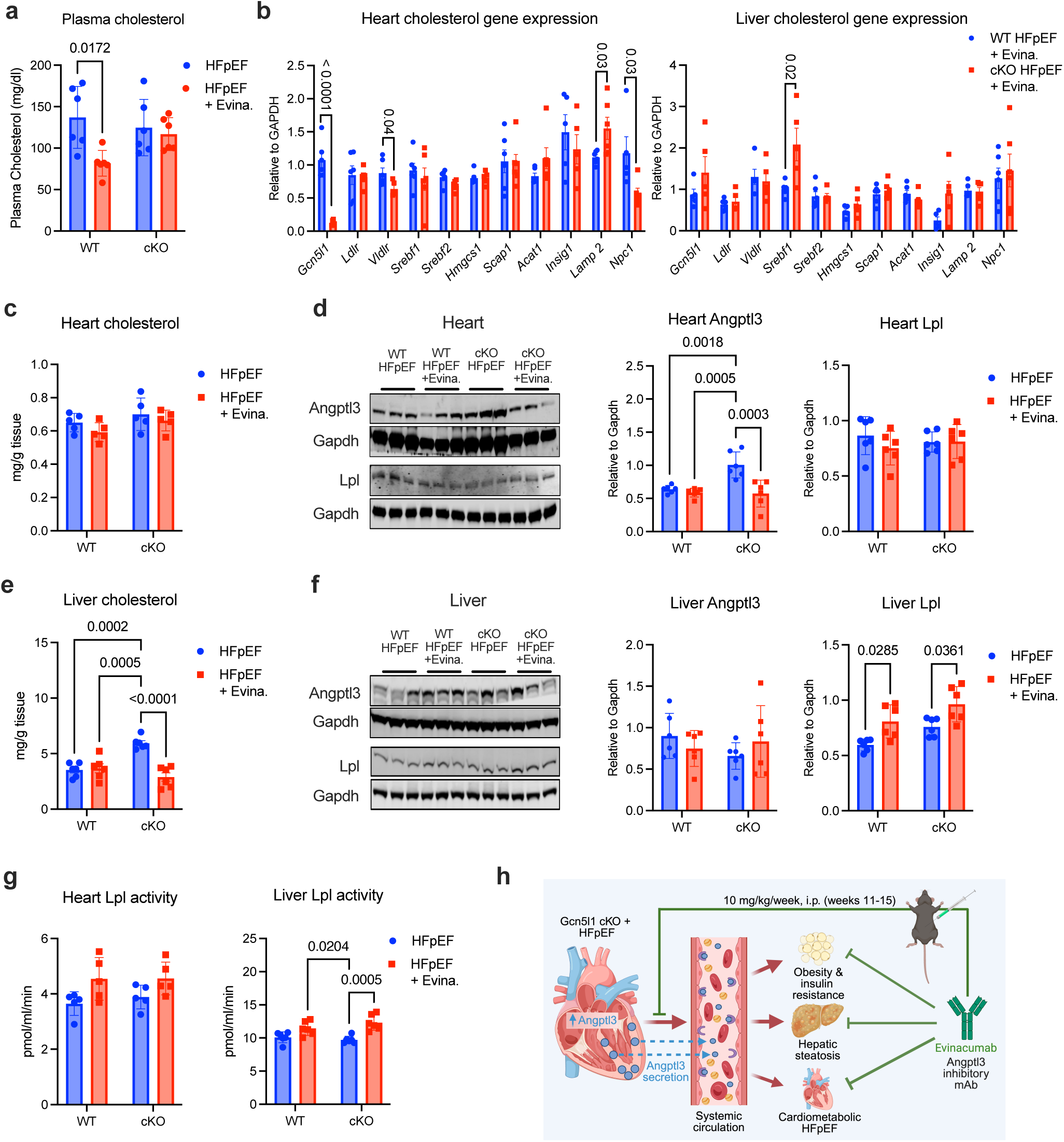
Rebalancing of the Angptl3-Lpl axis by Evinacumab. **a,** Biochemical measurement of plasma cholesterol in WT and Gcn5l1 cKO HFpEF mice treated with vehicle or Evinacumab. Data are mean ± SEM, with two-way ANOVA and Tukey’s post-hoc test, N = 6. **b,** Cholesterol metabolism gene expression in heart and liver of HFpEF mice. Data are mean ± SEM, with multiple unpaired two-tailed t-tests, N = 5. **c,** Biochemical measurement of heart cholesterol in WT and Gcn5l1 cKO HFpEF mice. Data are mean ± SEM, with two-way ANOVA and Tukey’s post-hoc test, N = 5. **d,** Protein abundance in HFpEF hearts after Evinacumab treatment. Data are mean ± SEM, with two-way ANOVA and Tukey’s post-hoc test, N = 6. **e,** Biochemical measurement of liver cholesterol in WT and Gcn5l1 cKO HFpEF mice. Data are mean ± SEM, with two-way ANOVA and Tukey’s post-hoc test, N = 6. **f,** Protein abundance in HFpEF livers after Evinacumab treatment. Data are mean ± SEM, with two-way ANOVA and Tukey’s post-hoc test, N = 6. **g,** Lpl activity in HFpEF hearts and livers after Evinacumab treatment. Data are mean ± SEM, with two-way ANOVA and Tukey’s post-hoc test, N = 5. **h,** Schematic of Evinacumab treatment in Gcn5l1 cKO HFpEF mice.

Conversely, we observed a significant decrease in cholesterol in Gcn5l1 cKO HFpEF livers (**Fig. 5e**), and no change in Angptl3 message or protein levels (**Fig. 5f** and **Supplementary Data Fig. 10b**). In contrast to the heart, Evinacumab treatment significantly increased both Lpl abundance and activity in Gcn5l1 cKO livers (**Fig. 5f,g**), suggesting elevated lipolysis activity in this tissue in response to Angptl3 inhibition. Combined, our findings imply that in the context of HFpEF, Evinacumab treatment helps to rewire cholesterol metabolism by shifting the balance away from Angptl3 expression, and towards increased Lpl abundance and activity (**Fig. 5h**), to aid lipolysis and promote lipid clearance.

## Discussion

Our study reveals a novel pathway linking cardiometabolic disease presentations in different tissues, driven by loss of Gcn5l1 in the heart, and a defect in systemic cholesterol metabolism. Despite a growing recognition of the heart as an endocrine signaling organ [5], there are very few examples of cardiac-derived factors that modulate systemic metabolism in the context of cardiometabolic disease. For example, natriuretic peptides released from the heart promote lipolysis and browning of adipocytes in human tissues and mice [19, 20], in addition to their canonical role in regulating blood pressure and volume. Cardiac expression of the Mediator complex member MED13 promotes leanness by upregulating energy metabolism in mouse liver and adipose tissue [21], with a similar finding in *Drosophila* being mediated by striated-muscle secretion of the Wingless peptide [22]. Finally, mitsugumin 53 (MG53) is released by the heart and skeletal muscle, which then binds to systemic insulin receptors to drive their degradation and promote insulin resistance [23]. The protective role of cardiomyocyte Gcn5l1 expression detailed in this report places it within this select group, and marks the identification of a new HFpEF-specific heart-to-liver signaling axis.

One unexpected finding of this study was the atypical tissue expression of several key lipid metabolism regulators (such as Angptl3 and Lpl) in mice and patients with HFpEF. Angptl3 is classically described as being almost exclusively expressed in the liver [13]. However, we detected its message and protein in Gcn5l1 cKO HFpEF mouse hearts, in human and rat cardiac cells with Gcn5l1 knockdown, and in human hearts from obese, type-2 diabetic patients (**Figs. 3,5** and **Supplementary Data Fig. 8a**). These data indicate that Gcn5l1 suppresses cardiac Angptl3 under normal conditions, and that this suppression is lost following Gcn5l1 depletion in the heart. Our findings here also support a recent large-scale screen of plasma samples from patients looking for novel proteins associated with heart failure development [24]. Using Mendelian randomization analysis and plasma proteomics, Shah and colleagues demonstrated that genetically-driven increases in circulating Angptl3 were significantly associated with increased left ventricle end diastolic volume and left atrial volume in humans – both echocardiographic markers of HFpEF-related diastolic dysfunction [24]. Combined with our findings reported here, these data suggest that Angptl3 may be a novel and underappreciated driver of cardiometabolic disease in the heart.

In direct contrast to Angptl3, Lpl is typically expressed in the heart, skeletal muscle, and adipose tissue, while very little is expressed in the adult liver [15]. Despite this, we found that *LPL* expression was highly upregulated in humans with MASLD (**Supplementary Data Fig. 8** and [16]), and that it was inversely correlated with Angptl3 expression in HFpEF mouse livers (**Figs. 3,5**). While hepatic Lpl has been shown to be upregulated in response to various stresses [16, 25], its function in healthy livers appears to be more important than its low gene expression level would otherwise suggest. For example, preclinical studies have shown that liver-specific deletion of Lpl results in a ∼25% decrease in total circulating levels of Lpl [26], while hepatic-specific Lpl overexpression prevents hypertriglyceridemia in rats with defective Lpl transport [27].

In terms of cardiac dysfunction, our finding that Lpl was decreased in WT HFpEF hearts matched a decrease in *LPL* message in myocardial biopsies from HFpEF patients (**Fig. 3f** and [14]). The potential causative nature of these observations has been corroborated in another preclinical HFpEF model where pharmacological blockade of Lpl, combined with cardiac-specific overexpression of Ldlr (to drive uptake of LDL cholesterol into the heart), leads to the recapitulation of several human HFpEF features in mice [28]. Combined with our current findings, these data suggest that there is a significant rewiring of heart and liver metabolism in HFpEF, which may include the upregulated expression of heart enzymes in the liver (and vice versa) to combat disease-related stress. In this context, the Angptl3-Lpl axis across heart and liver appears to be worthy of further investigation.

Elevated total cholesterol levels have long been associated with an increased incidence of heart failure with reduced ejection fraction (HFrEF), which is characterized by a loss of systolic function [29]. In contrast, higher cholesterol levels have received relatively less attention as a driver of HFpEF. However, recent observational studies have begun to show a potentially important role for controlling cholesterol levels in this cardiometabolic disease. The Netherlands-based PREVEND study, which has followed a cohort of >8500 patients for over 25 years, identified hypercholesterolemia as a top five risk factor for the future development of HFpEF in women [30]. In the United States, an observational study of ∼8000 patients demonstrated that newly-initiated statin use was associated with a ∼20% decrease in all-cause mortality and major cardiovascular events in recently diagnosed HFpEF patients after six years of tracking [31]. Importantly, these patients had no atherosclerotic cardiovascular disease at HFpEF diagnosis, suggesting the protective effects of statins were independent of coronary artery disease [31], a disorder typically linked to high cholesterol levels. In this context, the use of newer cholesterol-lowering drugs (such as the Angptl3 inhibitory monoclonal antibody Evinacumab; **Figs. 4,5**), may be part of a wider treatment regimen used to combat HFpEF in patients.

The function of Gcn5l1 as a regulator of cholesterol metabolism described here extends well beyond its previously hypothesized role in this pathway. Previous work has shown that Gcn5l1 is involved in the regulation of late endolysosomal processes, and as such can regulate LDL receptor recycling in hepatocytes [32]. Here, we show that multiple genes in cholesterol synthesis, cholesterol metabolism, apolipoprotein biology, and oxysterol-binding protein pathways are dysregulated in the heart by cardiomyocyte-restricted Gcn5l1 deletion, and that this leads to significant changes in cholesterol metabolism and abundance in the liver (**Fig. 2**). Furthermore, we identify Angptl3 as being specifically regulated by Gcn5l1 abundance in a number of *in vivo* and *in vitro* settings (**Figs. 4,5** and **Supplementary Data Fig. 7**), suggesting that the regulation of cholesterol metabolism by Gcn5l1 goes well beyond receptor recycling.

Finally, a number of patient mutations were recently described that result in reduced protein abundance of Gcn5l1 in the brain [33]. These mutations were associated with lysosomal defects and a loss of myelin production (hypomyelinating leukodystrophy), leading to the development of neurological disorders and early mortality [33]. As cholesterol is a major component of myelin, and circulating cholesterol cannot pass through the blood brain barrier [34], it is tempting to speculate that the reduced myelin observed in these patients was related to disrupted cholesterol synthesis in oligodendrocytes caused by mutation-driven Gcn5l1 depletion. Future studies will be required to understand the full impact of Gcn5l1 regulation on cholesterol metabolism in both single tissues, and on interorgan crosstalk pathways, across a range of human diseases.

## Acknowledgements

We would like to thank The Center for Organ Recovery and Education (CORE), the donors, and their families for their gifts to research.

## Author contributions

P.B. planned and executed experiments, analyzed raw data, and edited the manuscript. B.A.S.M., M.W.S., J.R.M., N.B., M.S-S., M.Z., A.V., R.G.R.M., C.L.H., and S.J.M. executed experiments. J.C.S. provided unique resources. S.K.R., B.A.K., S.S.S., S.L.G., and M.J.J. provided critical expertise and edited the manuscript. I.S. planned experiments, performed secondary analysis of data, and wrote the manuscript. All authors have reviewed the final manuscript and agree with the submission.

## Materials and correspondence

Requests for materials, along with any correspondence related to this article, should be addressed to Iain Scott, PhD.

## Supplementary materials

## Methods

### Generation of cardiomyocyte-specific Gcn5l1 knockout mice

All mice were bred and maintained in compliance with protocols approved by the University of Pittsburgh institutional animal care and use committee. Cardiomyocyte-specific conditional knockout mice on a C57BL/6J background were generated by crossing αMHC-MerCreMer transgenic mice (Jax B6.FVB (129)-Tg(Myh6-cre/Esr1*)1JMK/J) with mice harboring LoxP sites flanking exon 3 of the Gcn5l1 [35]. Gcn5l1^WT/WT;Cre+/-^ (WT) and Gcn5l1^FL/FL;Cre+/-^ (Gcn5l1 cKO) mice received a single intraperitoneal injection of tamoxifen at a dose of 40 mg/kg to induce Gcn5l1 deletion in floxed mice. Successful Gcn5l1 knockout was confirmed by qPCR and western blot analysis performed four weeks after tamoxifen administration. Both male and female mice were used for initial characterization (**Fig. 1** and **Supplementary Data Figs. 2,3**), with male mice being used for subsequent studies (**Figs. 2-5** and **Supplementary Data Figs. 4-10**).

### Two-hit model of heart failure with preserved ejection fraction (HFpEF)

A HFpEF phenotype was induced in both WT and Gcn5l1 cKO mice by administering a high-fat diet (HFD; Research Diets D12492; containing 60% calories from fat) and L-NAME (Sigma) at 0.5 g/L in drinking water [12]. The development of HFpEF was confirmed by the presence of increased body weight, insulin resistance, cardiac diastolic dysfunction (as shown by altered echocardiographic parameters e.g. E/e’ ratio, E/A ratio), left ventricular hypertrophy, and cardiac interstitial fibrosis. All experimental mice were euthanized according to approved protocols. Hearts, livers, skeletal muscle, and visceral adipose tissue were collected and snap-frozen for biochemical analysis. A subset of the hearts and livers from experimental animals were fixed in 4% paraformaldehyde solution for histological examination.

### Evinacumab intervention study

A HFpEF phenotype was induced in male WT and Gcn5l1 cKO animals as described above. Mice were randomly assigned to groups and administered either the Angptl3 inhibitory monoclonal antibody Evinacumab (MedChemExpress) at 10 mg/kg once weekly for four weeks, from the 11th to the 15th week of the study, or a vehicle control (sterile PBS). Insulin resistance and cardiac function in vehicle- and Evinacumab-treated mice were evaluated by intraperitoneal glucose tolerance test (IPGTT) and echocardiography. All experimental mice were euthanized according to approved protocols. Hearts, livers, skeletal muscle, and visceral adipose tissue were collected and snap-frozen for biochemical analysis. A subset of the hearts and livers from experimental animals were fixed in 4% paraformaldehyde solution for histological examination.

### Human tissue samples

Fresh human cardiac tissue samples were collected from the left ventricles of organ donors deemed not suitable for transplant, under a protocol approved by the University of Pittsburgh Committee for Oversight of Research and Clinical Training (CORID). Tissues were flash-frozen and stored at −80°C until processing. Study authors only had access to the de-identified tissue samples and their associated clinical phenotyping data, and no personally identifiable information was obtained.

### Intraperitoneal glucose tolerance test

Intraperitoneal glucose tolerance tests (IPGTTs) were performed on overnight-fasted mice. D-glucose was dissolved in sterile saline, and a glucose bolus (2 g/kg body weight) was injected intraperitoneally. Blood glucose levels were measured before glucose solution injection (*i.e.*, 0 min) and after 15, 30, 60, and 120 min. The data were analyzed, and an area under the curve calculated.

### Echocardiography

Cardiac function was evaluated by transthoracic echocardiography using a VisualSonics Vevo 3100 system equipped with an MS400 transducer (Visual Sonics Inc). Anesthesia was induced by 3% isoflurane and confirmed by lack of response to firm pressure on one of the hind feet. During echocardiogram acquisition, under body temperature was controlled using a heating pad, and isofluorane was reduced to 1.0-1.5% and adjusted to maintain heart rate in the range of 400-550 bpm. Left ventricular ejection fraction and other indices of systolic function were recorded from short-axis M-mode scans at the papillary muscle level under isoflurane-induced anesthesia in gently restrained mice. Apical 4-chamber views were used in anesthetized mice to evaluate diastolic function measurements using pulsed-wave and tissue Doppler at the level of the mitral valve. Global longitudinal strain (GLS) was recorded via parasternal long-axis view, and speckle tracking analysis was employed to confirm global longitudinal strain of the myocardium. Parameters collected include: heart rate (HR), left ventricular (LV) mass, left ventricular fractional shortening (LVFS), left ventricular ejection fraction (LVEF), peak Doppler blood inflow velocity across the mitral valve during early diastole (E), peak Doppler blood inflow velocity across the mitral valve during late diastole (A), isovolumic relaxation time (IVRT), and peak tissue Doppler of myocardial relaxation velocity at the mitral valve annulus during early diastole (e’). GLS was analyzed by speckle tracking analysis. At the end of the procedures, all mice recovered from anesthesia without difficulties. All parameters were measured at least 3 times, and averages are presented.

### High fat diet

For the high-fat diet comparison study, WT and Gcn5l1 cKO mice were provided with a high-fat diet (HFD; Research Diets D12492; containing 60% calories from fat) without L-NAME for the same timeframe as the HFpEF study (15 weeks).

### Histology

Heart and liver were harvested and fixed in 4% paraformaldehyde (PFA) overnight and processed for routine paraffin histology (5 μm sections stained with hematoxylin and eosin, and Masson’s trichrome). Cardiac fibrosis areas were visualized with a Leica DM2000 upright photomicroscope and quantified using ImageJ software version 2.0.

### Bulk RNA sequencing

Total RNA was isolated from the left ventricle of the heart using the RNeasy Plus Mini Kit (Qiagen, Cat. No. 74134). Approximately 2 μg RNA was used for bulk RNA sequencing performed by Azenta/GENEWIZ, based on company recommendations to identify differential gene expression patterns between the experimental groups. Raw data can be found here: <u>10.5281/zenodo.19484311</u>

### Proteomics

Mouse plasma samples were collected into anticoagulant-coated tubes and processed promptly by centrifugation to obtain plasma, which was aliquoted and stored at -80°C until analysis to minimize pre-analytical variability. For proteomic profiling, plasma proteins were depleted of abundant proteins using High-Select Top14 Abundant Protein Depletion Resin (Thermo Scientific). Protein digestion was carried out on 10 µg of protein from each sample according to the S-trap (Protifi) protocol. Samples were subject to reduction with 20 mM dithiothreitol, heated to 95°C for 10 min, cooled to room temperature, and alkylated by incubation with 40 mM iodoacetamide in darkness for 30 min at room temperature. Samples were centrifuged at 13000 x g for 8 min, acidified with 12% phosphoric acid at 1:10 concentration, and diluted six-fold in binding buffer containing 90% methanol and 100 mM TEAB at pH 7.1. Samples were dispensed onto S-trap columns which were washed with binding buffer and centrifuged at 4000 x g for 1 min. Columns were washed with binding buffer, centrifuged at 4000 x g for 1 minute, and transferred to clean tubes for incubation at 47°C in 50 mM TEAB for trypsin digestion at 1:10 enzyme/substrate. Samples were eluted with a series of solutions including 50 mM TEAB, 0.2% formic acid, and 50% acetonitrile with 0.2% formic acid, and underwent centrifugation at 1000 x g for 1 min after each eluent was added. Samples were then dried in a speedvac and resuspended in a solution of 3% acetonitrile and 0.1% TFA and desalted using Pierce Peptide Desalting Spin Columns (Thermo Scientific). Eluants were dried in a speedvac and resusupended in a solution of 3% acetonitrile and 0.1% formic acid to a final concentration of 0.2 µg/µL.

Mass spectrometry analysis was conducted on a Bruker timsTOF Pro2 coupled to a NanoElute 2. Approximately 200 ng of peptide was loaded onto an Aurora Ultimate C18 column (1.7 µm, 25 cm x 75 µm, IonOpticks) and eluted at 300 nL/min over a 60-minute gradient. The timsTOF Pro2 was set to PASEF scan mode and DDA with a scan range of 100-1700 *m/z* with 10 PASEF ramps. The TIMS was set to a 100 ms ramp and accumulation time (100% duty cycle) with a ramp rate of 9.43 Hz. Linear precursor repetitions were set to a target intensity of 20,000 and a target threshold of 2500 and active exclusion was set to 0.40 min. Collision energy was set to a base of 1.60 1/K_0_ [V-s/cm^2^] at 59 eV and 0.60 1/ K_0_ [V-s/cm^2^] at 20 eV. An isolation width was set to 2 *m/z* for <700 *m/z* and 3 *m/z* for >800 *m/z*.

Raw data files were analyzed with MSFragger v.4.1 in FragPipe v.22.0 and searched against the mouse SwissProt database (fragment mass tolerance = 20 ppm; precursor mass tolerance = 20 ppm; missed cleavages = 2; peptide length = 5 to 50; dynamic modifications: Oxidation +15.995 Da, Acetyl +42.011 Da; static modifications: Carbamidomethyl +57.021 Da). Peptide spectral matches were filtered using FDR = 1%. Raw data can be found here: <u>10.5281/zenodo.19484311</u>

### Metabolomics

Metabolic quenching and polar metabolite pool extraction was performed by adding ice cold 80% methanol (aqueous) at a ratio of 1:15 wt input tissue:vol. [^13^C]-creatinine, taurine-d_3_, lactate-d_3_ and alanine-d_3_ (Sigma-Aldrich) were added to the sample lysates as an internal standard for a final concentration of 10 μM. Samples were homogenized using an MP Bio FastPrep system using Matrix D (ceramic sphere) for 60 s at 60 Hz. The supernatant was then cleared of protein by centrifugation at 16,000 x g. Cleared supernatant (2 μL) was subjected to online liquid chromatography high resolution mass spectrometry (LC-HRMS) analysis. Briefly, samples were injected via a Thermo Vanquish UHPLC and separated over a reversed phase Thermo HyperCarb porous graphite column (2.1×100 mm, 3 µm particle size) maintained at 55 °C. For the 20 min LC gradient, the mobile phase consisted of the following: solvent A (water/0.1% FA) and solvent B (ACN/0.1% FA) at a flow rate of 0.3 mL/min. The gradient started at 1%B until 1 min where it increased to 15%B over the next 5 min. The organic increased to 98%B over the next 5 min and was held for 5 min before equilibration at initial conditions of 1%B for 5 min. The Thermo ID-X tribrid mass spectrometer was operated in both positive and negative ion mode, scanning in ddMS^2^ mode (2 μscans) from 70 to 800 *m/z* at 120,000 resolution with an AGC target of 2e^5^ for full scan, 2e^4^ for MS^2^ scans using HCD fragmentation at stepped (15, 35, 50) collision energies. Source ionization setting was 3.0 and 2.4 kV spray voltage, respectively, for positive and negative mode. Source gas parameters were 35 sheath gas, 12 auxiliary gas at 320 °C, and 8 sweep gas. Calibration was performed prior to analysis using the PierceTM FlexMix Ion Calibration Solutions (Thermo Fisher Scientific). Integrated peak areas were then extracted manually using Quan Browser (Thermo Fisher Xcalibur ver. 2.7). Untargeted differential comparisons were performed using Compound Discoverer 3.0 (Thermo Fisher) to generate a ranked list of significant compounds with tentative identifications from BioCyc, KEGG, and internal compound databases. Raw data can be found here: <u>10.5281/zenodo.19484311</u>

### Cholesterol quantification

Frozen mouse tissues (∼40 mg) were homogenized in 3 mL of chloroform:methanol (2:1) mixture in glass vials and vortexed thoroughly. The homogenate was incubated for 60-90 min at room temperature while rotating on a shaker with intermittent vortexing. The homogenate was then acidified with 1 mol/L H_2_SO_4_, and centrifuged at 1000 rpm for 10 min at room temperature. The lipid fractions in the lower organic phase were collected and transferred to clean glass vials. A small fraction (20-30 μL) of lipid fraction was transferred to a new glass vial and evaporated completely. Lipid levels were determined using colorimetric Infinity Cholesterol Reagent (Thermo Fisher Scientific) and normalized to organ weight.

### Thin layer chromatography (TLC)

Hepatic lipids were extracted from mouse livers and dried completely under a N-EVAP nitrogen evaporator (Organomation), then resuspended with 100 μL chloroform:methanol mixture (2:1). Before running the TLC plate (EMD Millipore), the running chamber was equilibrated with 200 ml of the solvent system containing petroleum ether:ethyl ether:acetic acid mixture (25:5:1). The TLC plate was baked at 75°C for 30 min, and 25 μL aliquots of resuspended lipid extract and lipid standard (Nu-Chek-Prep) were loaded onto the TLC plates. Lipids were separated using the solvent system. The plate was then transferred to an equilibrated iodine tank for 30-45 min for staining, and images were captured with CanoScan LidE 220 imager (Canon).

### LPL Activity assay

LPL activity of heart and liver tissues were assessed using a commercially available Lipoprotein Lipase Activity Assay Kit (Fluorometric) (AbCam, Cat. No. ab204721) following the manufacturer’s instructions.

### Generation of GCN5L1 KD stable cell lines

Human cardiac AC16 cells (Millipore) were transduced with lentiviral particles at a multiplicity of infection (MOI) of 10 with scrambled control or GCN5L1 shRNA (Sigma-Aldrich, USA), followed by puromycin selection. GCN5L1 KD was confirmed by RT-qPCR and western blotting. Control and GCN5L1 KD cells were washed with 1X PBS and lysed in CHAPS lysis buffer with protease and phosphatase inhibitors.

### Isolation of NRCMs and GCN5L1 knockdown

Hearts were quickly excised from one to three-day-old Sprague Dawley neonates, and cardiac myocyte isolation was accomplished by digestion with 0.04% trypsin and type II collagenase (0.4mg/mL; Worthington, Cat. No. LS004176) in Krebs-Henseleit bicarbonate buffer at 37 °C. Non-cardiomyocyte cells were removed by rapid attachment (90 mins incubation in culture dishes). Cardiomyocytes were plated at a density of 2 × 10^5/mL in DMEM containing 10% FBS and 0.1 mM BrdU to stop the growth of fibroblasts. Twenty-four hours after cell plating, the medium was changed to serum-free DMEM containing 0.1% insulin-transferrin-selenium (Thermo Fisher Scientific Cat No. 41400045). NRCMs were transfected with scrambled control siRNA or GCN5L1 siRNA (Origene, Cat No. SR15522) using Lipofectamine RNA iMAX Transfection Reagent (Thermo Fisher Scientific) according to the manufacturer’s protocol.

### LPL Activity assay in AC16 cells

Control and GCN5L1 KD AC16 cells were cultured in a 10 cm dish for 24 h. Cell pellets were collected by trypsinization and washed using PBS three times, then cells were lysed in ice-cold 1X PBS using a Dounce homogenizer. LPL activity was assessed as described above.

### RT-qPCR

RNA was extracted from heart, liver, skeletal muscle, visceral adipose tissue, AC16 cells, and NRCMs using the RNeasy kit (Qiagen), with a μLITE analyzer (BioDrop) used to evaluate the quality and quantity. RNA (1 μg) was used to synthesize cDNA using the Maxima Reverse Transcriptase kit (Thermo Fisher Scientific,), and gene expression studies were performed with mouse specific primers for Gcn5l1 (Cat. No. QT00016002), Ldlr1 (Cat. No. QT00109823), Vldlr1 (Cat. No. QT00101360), Srebf2 (Cat. No. QT01045870), Scap1 (Cat. No. QT00167811), Acat1 (Cat. No. QT00166684), Insig1 (Cat. No. QT00120120), Lamp2 (Cat. No. QT00101059), Npc1 (Cat. No. QT00106400), Hmgcs2 (Cat. No. QT00169029), Hmgcs1 (Cat. No. QT00132461), and Gapdh (Cat. No. QT01658692); or with human specific primers for GCN5L1(Cat. No. QT01063069), ANGPTL3 (Cat. No. QT00034377), and GAPDH (Cat. No. QT01658692); or with rat specific primers for GCN5L1 (Cat. No. QT01594159), ANGPTL3 (Cat. No. QT02550233), and GAPDH (Cat. No. QT00132461). All primers and SYBR green master mix (Cat No. QT01082004) were obtained from Qiagen.

### Immunoblot analysis

Protein extracts from frozen mouse heart, liver, skeletal muscle, and visceral adipose tissue were prepared by lysis in ice-cold CHAPS lysis buffer (150 mM NaCl, 50 mM Tris HCL pH 7.4, 1% Triton-X 100, 0.5% sodium deoxycholate, 0.1% SDS, 5 mM EDTA, 2 mM EDTA) containing protease and phosphatase inhibitors. Protein extracts from control and GCN5L1 KD cells were obtained by lysis in ice-cold CHAPS lysis buffer (150 mM NaCl, 50 mM Tris HCL pH 7.4, 1% Triton-X 100, 0.5% sodium deoxycholate, 0.1% SDS, 5 mM EDTA, 2 mM EDTA) containing protease and phosphatase inhibitors. Proteins were separated by SDS-PAGE on 4-12% or 12% gradient gels (Invitrogen), and transferred to nitrocellulose membranes. An Odyssey scanner (LI-COR version 3.0) was used as a detection system. Protein expression was measured using a fluorescently labelled secondary antibody, and protein band intensities were analyzed with ImageJ software. Proteins were detected with the following primary antibodies from Protein Tech: GAPDH, Cat No: 60004-1-1g; 1:5000, ANGPTL3: 86193-1-RR 1:2000, LPL Cat No: 28602-1-AP; 1:3000, LDLR: Cat No 10785-1-AP; 1:4000, NPC1: 13926-1-AP; 1:1000. GCN5L1 antibody was custom produced by Covance and validated here [1].

### Statistical analysis

All data are presented as means ± SEM. Statistical analyses of studies with two variables were conducted using two-way ANOVA followed by Tukey’s post hoc test. Studies involving a single variable were evaluated with unpaired, two-tailed Student’s t tests. A *P* value ≤ 0.05 was considered significant, and all analyses were performed in GraphPad Prism 10.0. All experiments were performed with at least three biological replicates. No statistical analysis was used to predetermine sample sizes; estimates were made based on our previous experience, experimental approach, animal availability, and feasibility required to obtain statistically significant results. Experimental animals were randomly assigned to each group. Investigators were blinded to the genotypes of the individual animals during both the experiments and the experimental analysis.

## Supplementary Data Figures

**Supplementary Data Fig. 1:**
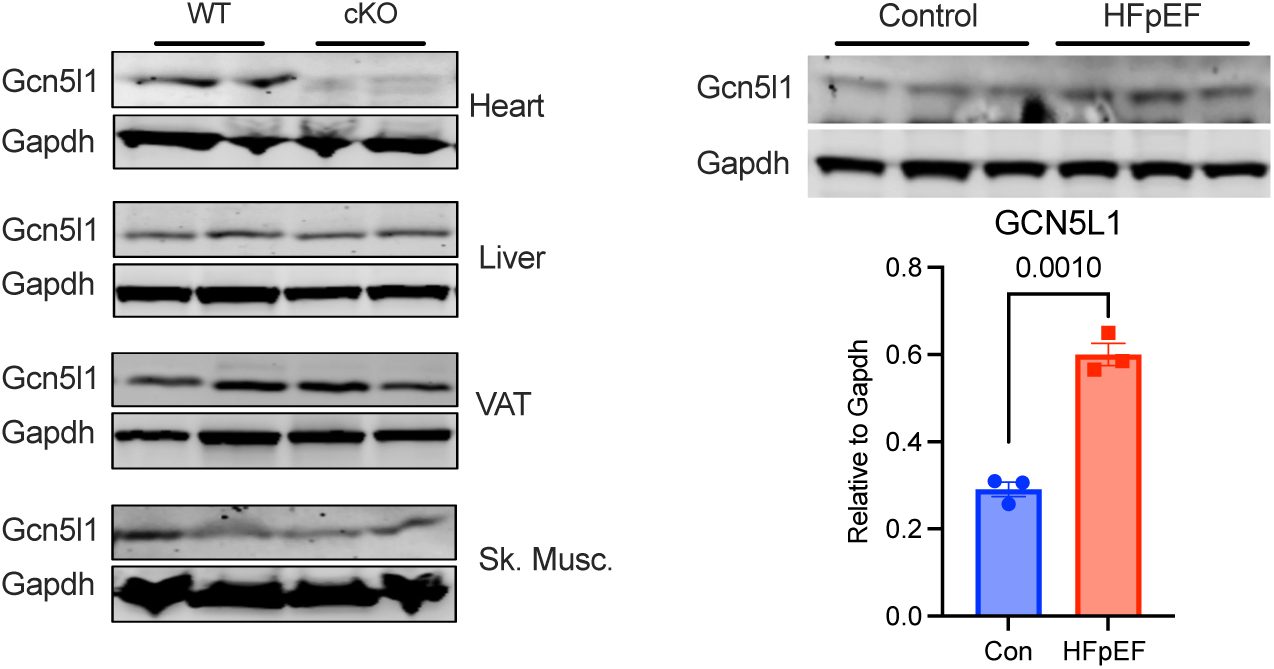
Gcn5l1 expression in WT and Gcn5l1 cKO mice. **a,** Gcn5l1 protein abundance in heart, liver, visceral adipose tissue (VAT) and skeletal muscle (Sk. Musc.). **b,** Gcn5l1 protein abundance in WT Control and HFpEF hearts. Data are mean ± SEM, with two-tailed t-test. N = 3.

**Supplementary Data Fig. 2:**
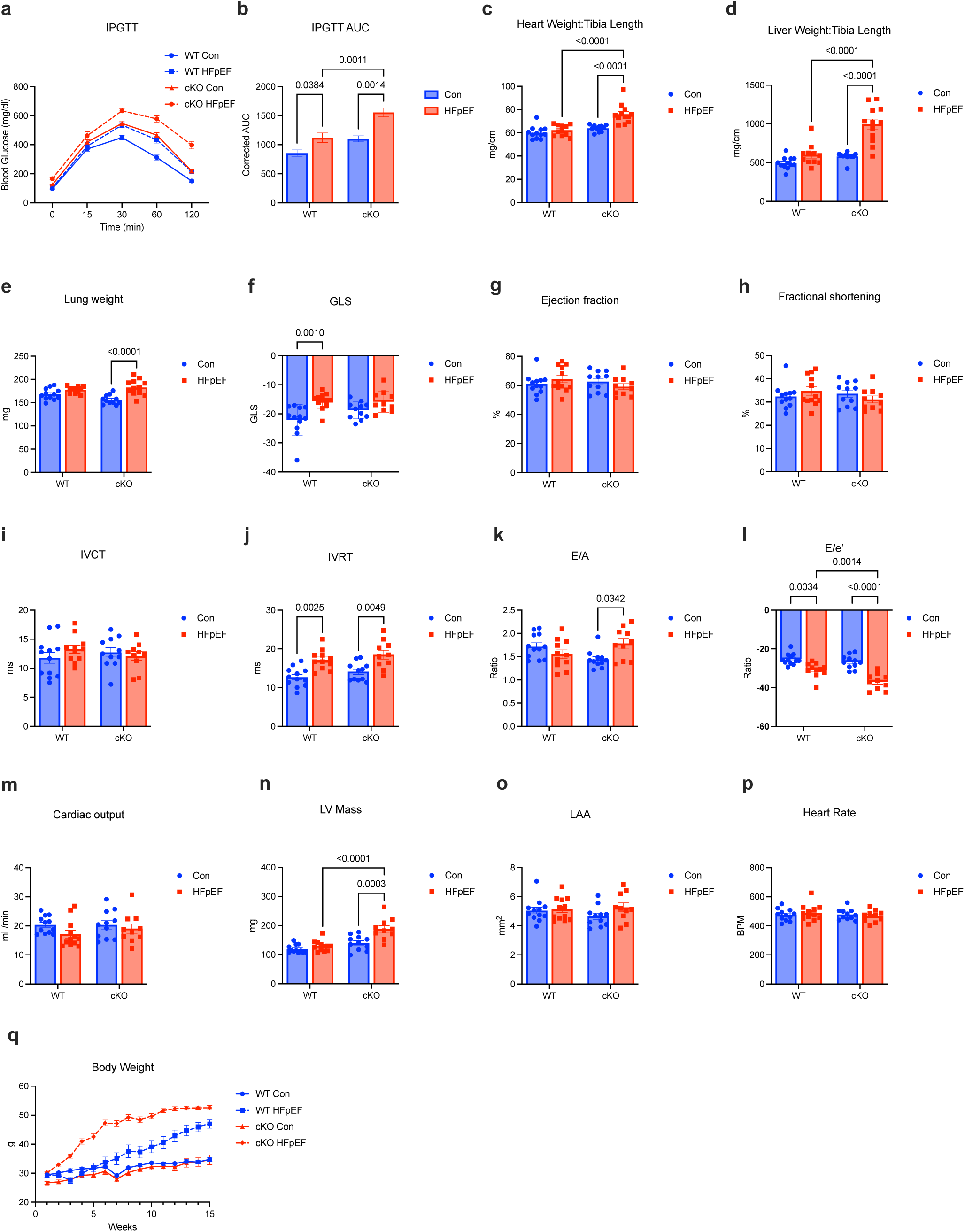
Metabolic and cardiac phenotyping in male mice. **a,b,** IPGTT curves and AUC. Data are mean ± SEM, with two-ANOVA and Tukey’s post-hoc test, N = 8. **c-p,** Full echocardiography analysis of cardiac function. Data are mean ± SEM, with two-ANOVA and Tukey’s post-hoc test, N = 11 (WT) or 12 (cKO). **q,** Body weight curves over course of study, N = 8. Some of the data presented in this figure are repeated from Fig. 1 of the main manuscript for ease of comparison.

**Supplementary Data Fig. 3:**
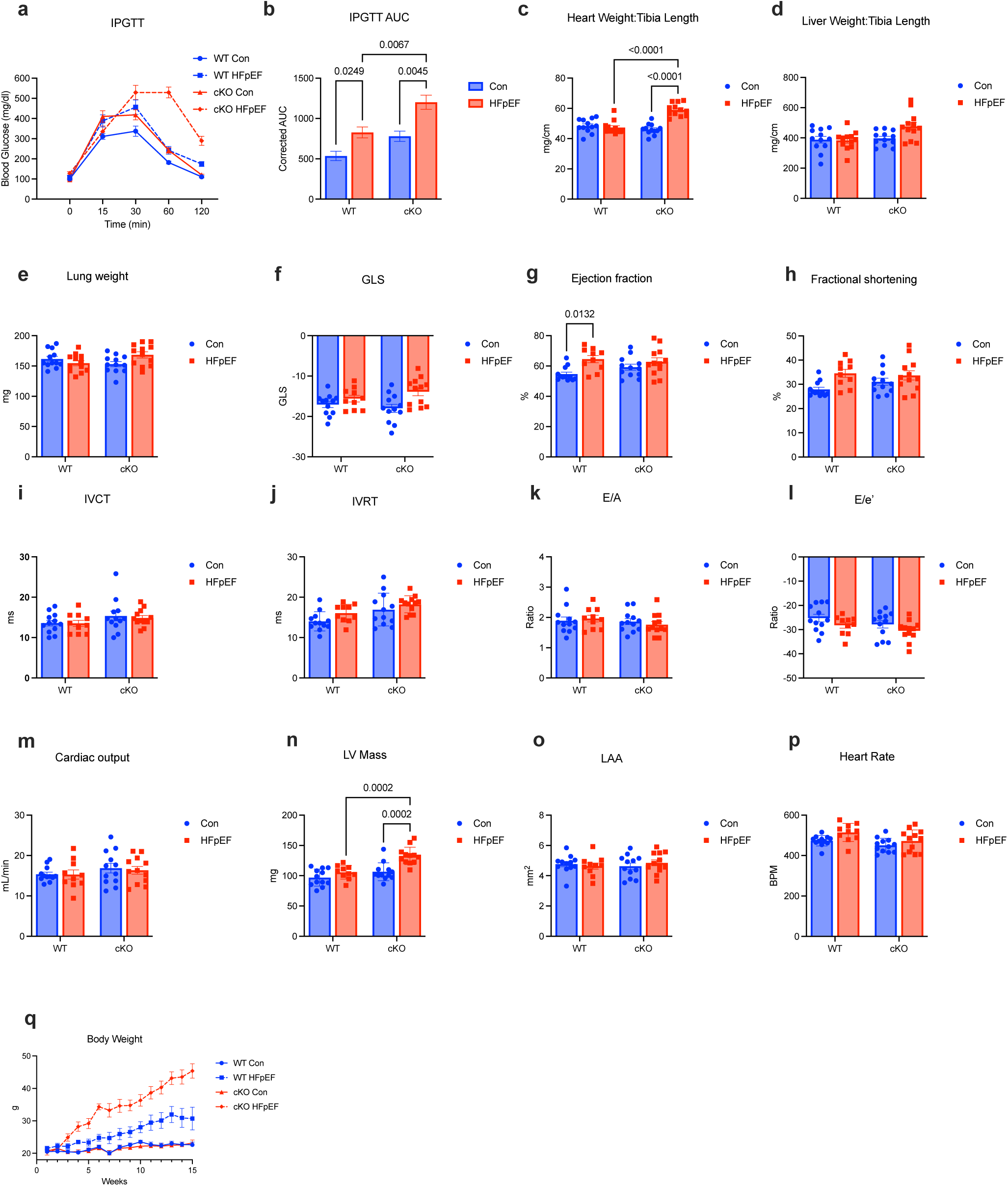
Metabolic and cardiac phenotyping in female mice. **a, b,** IPGTT curves and AUC. Data are mean ± SEM, with two-ANOVA and Tukey’s post-hoc test, N = 8. **c-p,** Full echocardiography analysis of cardiac function. Data are mean ± SEM, with two-ANOVA and Tukey’s post-hoc test, N = 11 (WT) or 12 (cKO). **q,** Body weight curves over course of study, N = 8.

**Supplementary Data Fig. 4:**
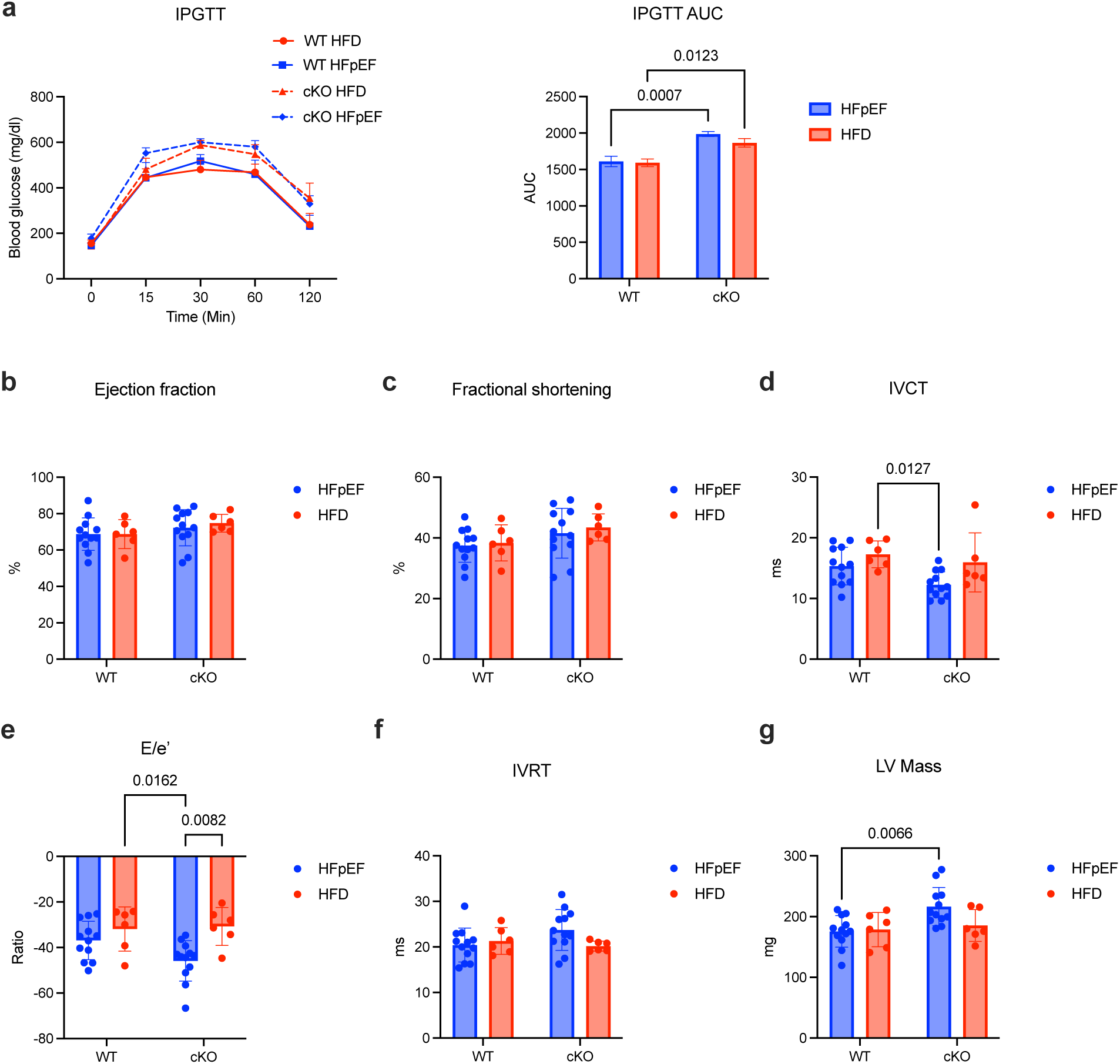
Metabolic and cardiac phenotyping in HFD vs. HFpEF mice. Echocardiography analysis of cardiac function. Data are mean ± SEM, with two-ANOVA and Tukey’s post-hoc test, N = 12 (HFpEF) or 6 (HFD).

**Supplementary Data Fig. 5:**
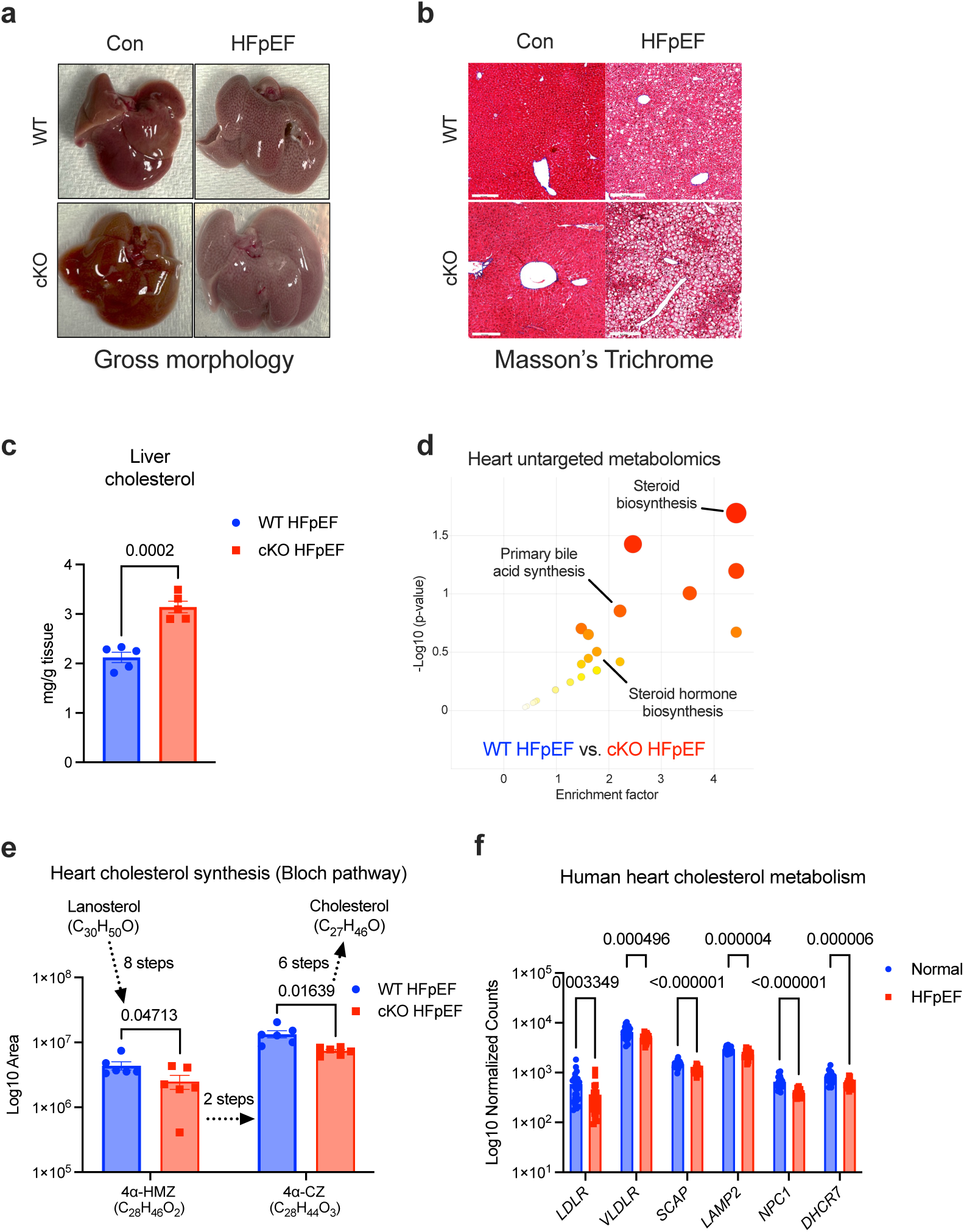
Liver phenotyping and cardiac cholesterol metabolism. **a,b,** Gross liver morphology and Masson’s trichrome staining of mouse livers. Scale bar = 200 μm. **c,** Biochemical analysis of total liver cholesterol in HFpEF mouse livers. Data are mean ± SEM, with a two-tailed t-test, N = 5. **d,** Untargeted heart metabolomics from WT and Gcn5l1 cKO HFpEF hearts. N = 8. **e,** Bloch pathway metabolite analysis from HFpEF mouse hearts. Data are mean ± SEM, with multiple two-tailed t-tests, N = 8. **f,** Heart cholesterol metabolism gene expression from a published human HFpEF data set [2]. Data are mean ± SEM, with multiple unpaired two-tailed t-tests, N = 25 (Normal) or 28 (HFpEF).

**Supplementary Data Fig. 6:**
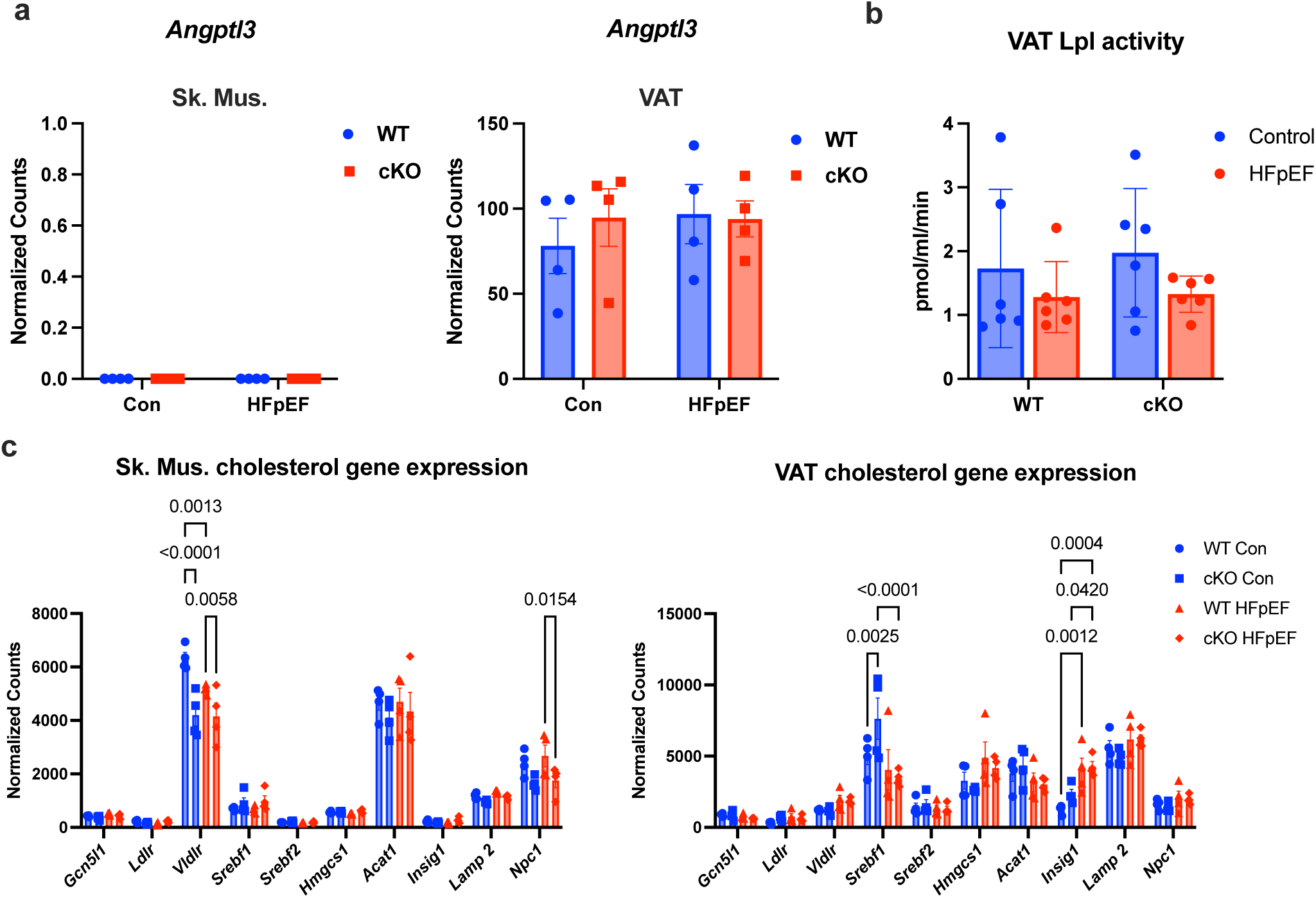
Cholesterol regulation in skeletal muscle and visceral adipose tissue. **a,** Gene expression from bulk RNA-seq. Data are mean ± SEM, with two-way ANOVA and Tukey’s post-hoc test, N = 4. **b,** Lpl activity in VAT. Data are mean ± SEM, with two-way ANOVA and Tukey’s post-hoc test, N = 6. **c,** Cholesterol gene expression from bulk RNA-seq. Data are mean ± SEM, with two-way ANOVA and Tukey’s post-hoc test, N = 4.

**Supplementary Data Fig. 7:**
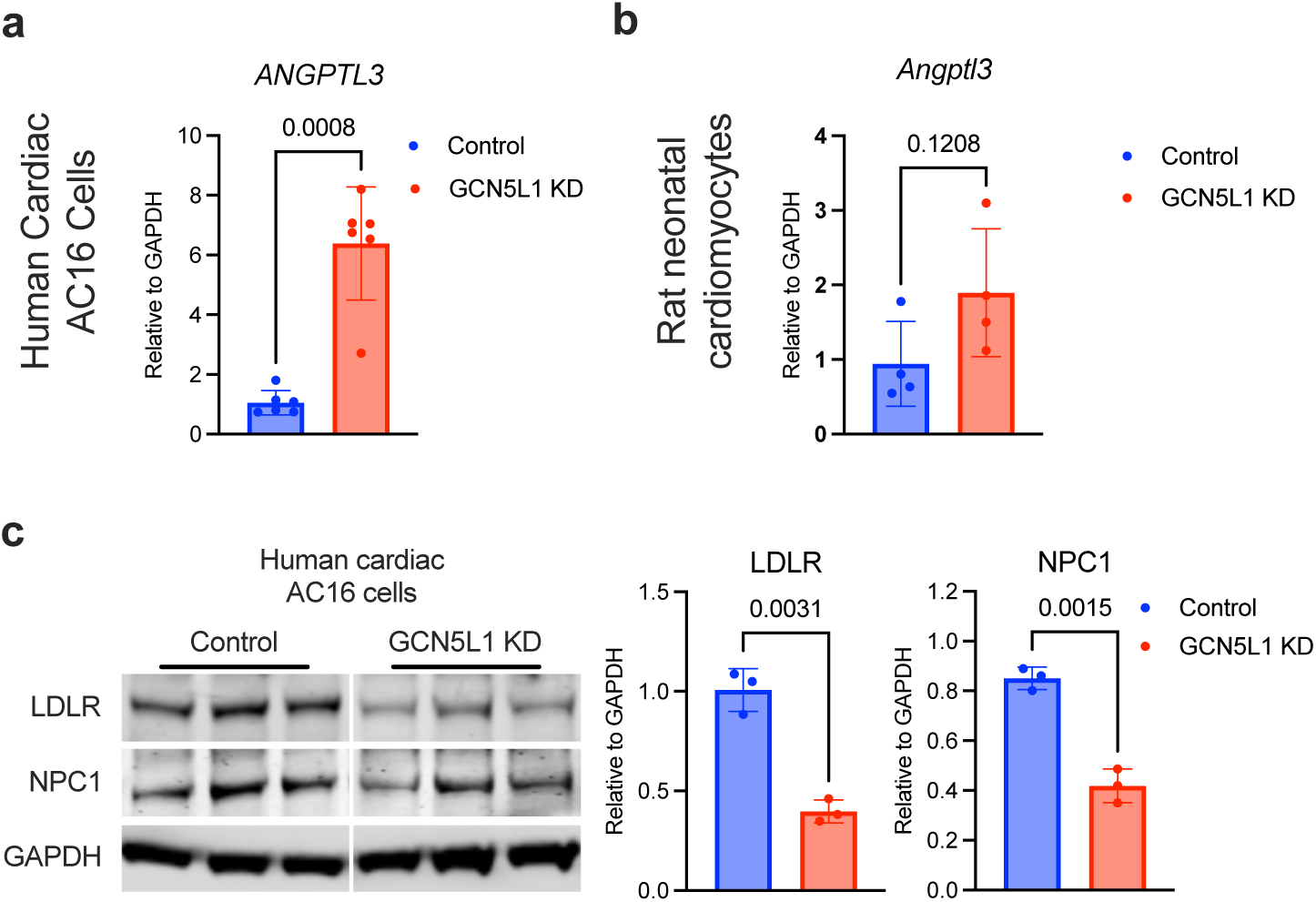
*In vitro* modeling of Gcn5l1 depletion in cardiac cells. **a,b,** Angptl3 gene expression measured by qPCR in human cardiac AC16 cells and rat primary neonatal cardiomyocytes following Gcn5l1 knockdown. Data are mean ± SEM, with two-tailed t-test, N = 6 (AC16) or 4 (Rat neonatal cardiomyocytes). **c,** LDLR and NPC1 protein abundance in human cardiac AC16 cells following GCN5L1 knockdown. Data are mean ± SEM, with two-tailed t-test. N = 3.

**Supplementary Data Fig. 8:**
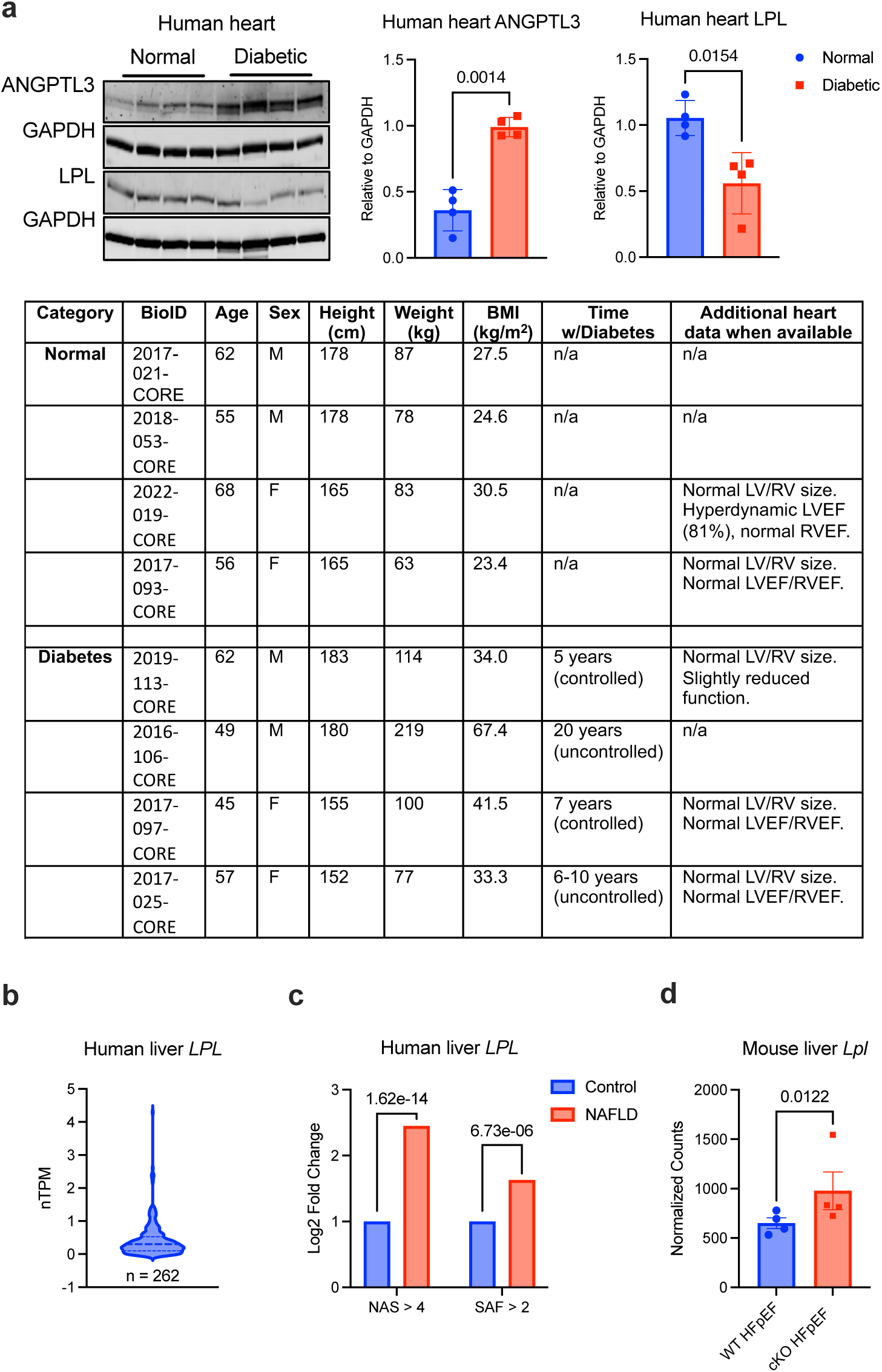
ANGPTL3 and LPL expression in diabetes and NAFLD. **a,** Protein abundance of ANGPTL3 and LPL in hearts from normal and diabetic individuals. Data are mean ± SEM, with two-tailed t-test, N = 4. **b,** *LPL* gene expression in 262 normal human livers. Data were obtained from The Human Protein Tissue Atlas (https://www.proteinatlas.org/ENSG00000175445-LPL/tissue/liver). **c,** *LPL* gene expression from a published human MASLD/NAFLD data set [3]. NAS = NAFLD activity score, SAF = FLIP steatosis, activity, and fibrosis score. Scores above 4 and 2, respectively, denote advanced MASLD progression. Data are mean ± SEM, with unpaired two-tailed t-test, N = 22 (Control) and 206 (NAFLD). **d,** *Lpl* gene expression in HFpEF mouse livers. Data are mean ± SEM, with two-tailed t-test, N = 4.

**Supplementary Data Fig. 9:**
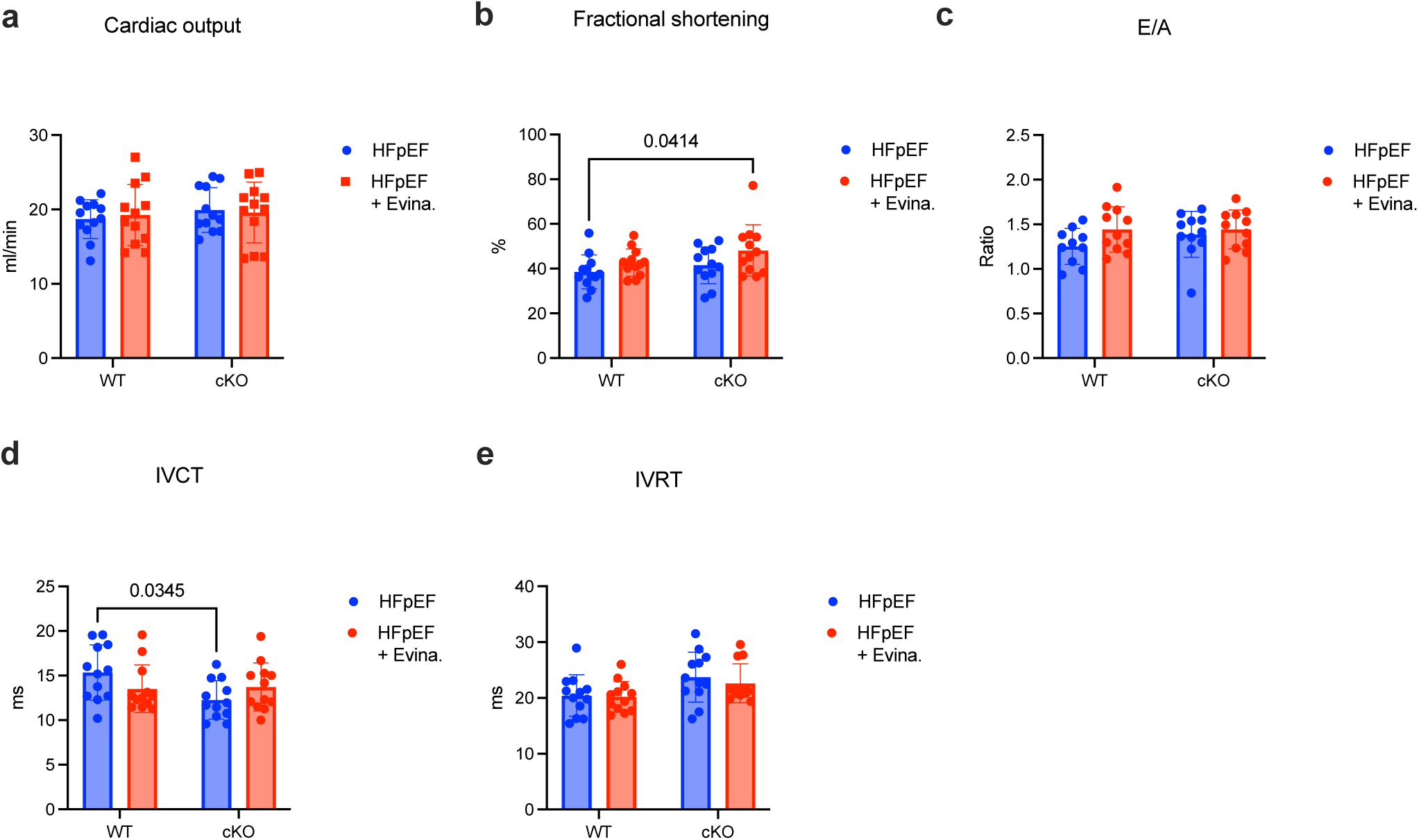
Cardiac phenotyping after Evinacumab treatment. Echocardiography analysis of cardiac function. Data are mean ± SEM, with two-ANOVA and Tukey’s post-hoc test, N = 12.

**Supplementary Data Fig. 10:**
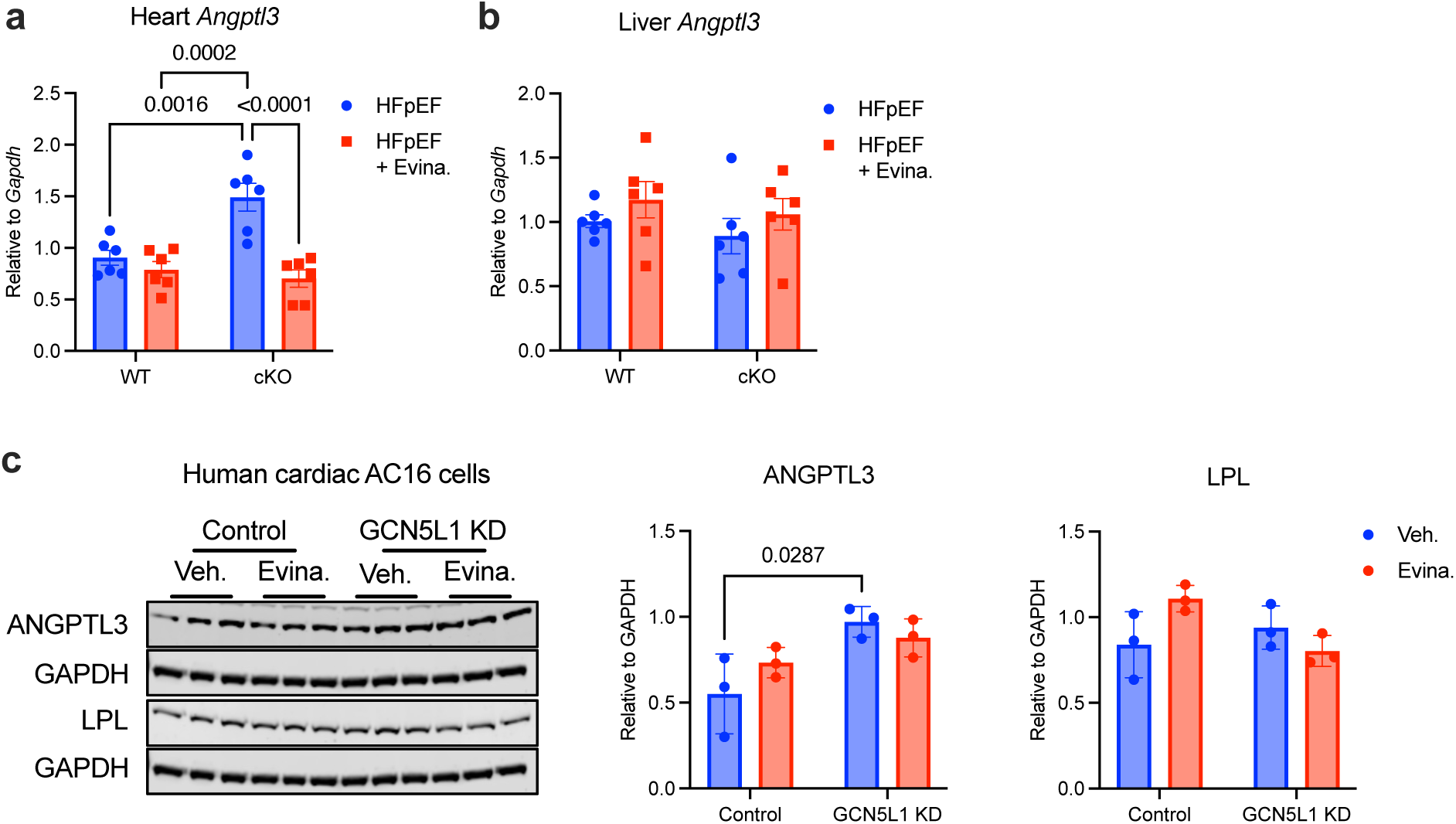
Angptl3 and Lpl expression after Evinacumab treatment. **a,b,** Heart and liver *Angptl3* gene expression measured by qPCR. Data are mean ± SEM, with two-way ANOVA and Tukey’s post-hoc test, N = 6. **c,** Protein abundance in Control and GCN5L1 knockdown cells treated with vehicle (PBS) or Evinacumab (10 μg/mL) for 24 h. Data are mean ± SEM, with two-way ANOVA and Tukey’s post-hoc test, N = 3.

